# A targeted mass-spectrometry plasma proteome for simultaneous multi-disease screening

**DOI:** 10.64898/2026.07.18.739311

**Authors:** Ahrum Son, Jaeho Ji, Eunjeong Han, Yunho Choi, Jumin Park, Heejin Lee, Sujin Park, Hyunsoo Kim

**Author notes:** Correspondence: Hyunsoo Kim – Department of Convergent Bioscience and Informatics, Chungnam National University, Daejeon 34134, Republic of Korea. Ahrum Son – Graduate School of Medical Science, University of Ulsan, Ulsan 44610, Republic of Korea.

## Abstract

Population health screening is fragmented across dozens of single-condition assays. We asked whether one targeted mass-spectrometry measurement of the plasma proteome could screen simultaneously for many common diseases. Using multiple-reaction-monitoring LC-MS/MS, we quantified 126 proteins (509 peptides) directly in crude plasma from 490 individuals spanning 16 diseases (eight cancers, three inflammatory or renal conditions, and five metabolic and lifestyle disorders) plus age- and sex-matched controls, each protein referenced to a stable-isotope-labelled internal standard. Proteome variation was organised primarily by disease rather than demographics. Cross-validated classifiers detected each disease against matched controls with a mean AUROC of 0.979, a 17-way model identified the specific disease at roughly ten times chance, and a 20-protein subset reproduced full-panel performance. We show that these within-cohort estimates are optimistic upper bounds because each disease was acquired as a separate case-control batch without independent validation. We quantify this, establishing analytical feasibility and defining the validation required for translation.

## Introduction

Early detection remains the most effective lever on outcomes for most common diseases^1^, yet the infrastructure of population screening is built one condition at a time. A routine health examination bundles a fasting glucose, a lipid panel, liver enzymes, a handful of tumor markers, and imaging— each a separate assay with its own sample handling, reference range, turnaround, and failure modes. This fragmentation is costly and incomplete: it multiplies pre-analytical variation, leaves gaps between the conditions that any single test was designed to catch, and scales poorly to the asymptomatic general population in whom early detection would be most valuable and the classical criteria for a screening program are hardest to satisfy^2^. A single measurement that simultaneously reported on many organ systems would change the economics of screening, provided it were quantitative, reproducible, and inexpensive enough to deploy at scale.

The plasma proteome is, in principle, an ideal substrate for such a measurement. Blood perfuses every tissue, and the proteins it carries encode a broad, real-time picture of physiological state—a phenotypic readout that, unlike the genome, reflects current status rather than fixed probabilistic risk^3,4^. Large population surveys have shown that plasma protein levels systematically track age, genetic variation, and disease across thousands of individuals^5, 6, 7, 8^. Two obstacles, however, have kept plasma proteomics from a general screening role. First, plasma protein abundance spans more than ten orders of magnitude, and the medically informative proteins are frequently among the least abundant^9^. Untargeted discovery platforms, which sample the proteome stochastically, struggle to quantify these low-abundance species reproducibly and consistently across large cohorts^3^. Second, and more consequentially, the path from a discovered protein signature to a deployable clinical test is notoriously unreliable: the great majority of candidate biomarkers fail to reproduce in independent cohorts, and only a handful of new protein markers reach clinical use each year^10, 11, 12, 13^. The dominant causes of this attrition—insufficient analytical reproducibility, poor cross-laboratory transferability, and confounding introduced by study design—are properties of the measurement and the cohort rather than of the underlying biology^10, 14^.

Targeted mass spectrometry in multiple-reaction-monitoring (MRM) mode addresses the analytical half of this problem directly. Rather than sampling the proteome at random, an MRM assay monitors a predefined set of peptide transitions, each quantified against a spiked stable-isotope-labelled (SIL) internal standard^15, 16^. This design delivers absolute, internal-standard-referenced quantitation that is reproducible within and between laboratories and linear across a wide abundance range^17, 18, 19, 20^, and it is already the analytical basis of clinical mass-spectrometry testing performed in accredited laboratories^21, 22^. Careful empirical selection of signature peptides and transitions underpins the specificity of such assays^23^. Because each target is defined in advance, an MRM panel can be multiplexed to scores or hundreds of proteins in a single injection^24, 25^ without the antibody-development and cross-reactivity limitations that constrain multiplexed affinity platforms^26, 27^. Targeted proteomics has accordingly been used to build reproducible, cancer-associated protein assays in body fluids^28^ and disease-specific, blood-based classifiers and liquid-biopsy signatures^29, 30^. Efforts to read many conditions from a single blood draw have nonetheless been dominated by nucleic-acid approaches and have focused almost exclusively on cancer. Multi-analyte tests based on circulating tumor DNA, on cell-free-DNA fragmentation or methylation, or on combined DNA and protein panels can detect and in some cases localize several malignancies from plasma^31, 32, 33, 34^, and targeted-methylation assays now screen for dozens of cancers at high specificity^35^. But somatic-mutation and methylation content is shared across tumur types and is largely uninformative for the substantial burden of common non-malignant disease—the metabolic, inflammatory, and renal conditions that dominate routine health screening. A protein assay that spanned cancers alongside these common non-malignant conditions, built on targets already validated as clinical or laboratory-developed-test (LDT) markers, would occupy a distinct and underexplored niche^36, 37^.

Here we test whether a single MRM assay, designed around proteins with existing clinical or LDT relevance and measured directly in crude plasma, can serve as a general multi-disease screen. We profiled 490 individuals spanning 16 diseases—eight cancers, three inflammatory or renal conditions, and five metabolic and lifestyle disorders—together with age- and sex-matched normal controls, quantifying 126 proteins against SIL internal standards. We then evaluate three questions in turn: whether the proteome separates disease from health; whether individual diseases can be detected against matched controls; and whether the specific disease can be identified. Throughout, we treat the central threat to any screening claim—that apparent discrimination reflects cohort and batch structure rather than biology—not as an assumption to be set aside, but as a quantity to be measured, and we bound its contribution explicitly^14, 38^.

## Methods

### Study cohort and clinical metadata

Plasma samples were obtained from 490 participants distributed across 17 groups: a normal-control group (NC, n = 30) and 16 disease groups. The disease groups comprised eight cancers—breast, colon, thyroid, pancreatic, liver, bladder, kidney, and prostate (n = 30 each); three inflammatory or renal conditions—glomerulonephritis (n = 10), cholecystitis (n = 30), and pancreatitis (n = 30); and five metabolic or lifestyle disorders—hyperlipidemia, hypertension, type-1 diabetes, type-2 diabetes, and obesity (n = 30 each).

Clinical metadata were harmonized directly from the source clinical records for each group. Because the 17 groups were documented in heterogeneous record formats, harmonisation required reconciling variable column names and encodings across sheets: age fields (recorded variously as first-participation age, current age, or intake age), sex fields (recorded as M/F or as numeric codes depending on group), and laboratory fields with inconsistent naming (for example, triglycerides recorded as either “TG” or “Triglyceride”). Height, weight, systolic and diastolic blood pressure, and routine chemistry and haematology values were harmonized into a common schema, and body-mass index was recomputed from height and weight where not directly recorded. Two data-entry values were corrected during harmonization: one type-2-diabetes participant carried an implausible recorded age of 168 years and was set to missing, and one hypertension participant with a recorded age of 93 years was retained as plausible. After harmonization, the normal-control group had a mean age of 52.4 ± 6.6 years (range 43–73), balanced sex (15 men, 15 women), and mean BMI 24.2 kg/m²— demographically representative of, rather than younger than, the disease groups. Full per-group demographics and clinical chemistry are in **Table 1, Table S1, and Fig. S1**.

**Table 1.** Demographic and clinical characteristics of the study cohort.

| Disease | SuperClass | N | Male_n | Female_n | Male_pct | Age_mean | Age_sd | BMI_mean | BMI_sd | SBP_mean | Glu_mean | Chol_mean | Age vs<br>NC<br>(p-value) | BMI vs<br>NC<br>(p-value) | Sex vs<br>NC<br>(p-value) |
| --- | --- | --- | --- | --- | --- | --- | --- | --- | --- | --- | --- | --- | --- | --- | --- |
| Normal control | Normal | 30 | 15 | 15 | 50 | 52.4 | 6.6 | 24.2 | 2.7 | 77.6 | 91.2 | 168.2 | - | - | - |
| Breast cancer | Cancer | 30 | 0 | 30 | 0 | 61.4 | 15.5 | 25.5 | 4 | 76.3 | 131.3 | 173.2 | 0.0516 | 0.1537 | 0.0000 |
| Colon cancer | Cancer | 30 | 18 | 12 | 60 | 67.6 | 9.9 | 23.3 | 2.2 | 78.5 | 122.5 | 167.2 | 0.0000 | 0.1494 | 0.6042 |
| Thyroid cancer | Cancer | 30 | 9 | 21 | 30 | 51.5 | 15 | 25.7 | 5.2 | 77.5 | 108.3 | 206.4 | 0.8474 | 0.3871 | 0.1872 |
| Pancreatic cancer | Cancer | 30 | 13 | 17 | 43.3 | 68.1 | 9.6 | 23 | 3.6 | 75.3 | 152.1 | 134.9 | 0.0000 | 0.1154 | 0.7961 |
| Liver cancer | Cancer | 30 | 24 | 6 | 80 | 64.4 | 8.8 | 24.6 | 4.1 | 76.4 | 136.9 | 167.2 | 0.0000 | 0.9823 | 0.0292 |
| Bladder cancer | Cancer | 30 | 26 | 4 | 86.7 | 71 | 12.6 | 25.2 | 3.7 | 78.9 | 130.7 | 164 | 0.0000 | 0.3403 | 0.0048 |
| Kidney cancer | Cancer | 30 | 22 | 8 | 73.3 | 65.2 | 11.6 | 25.1 | 3.4 | 80.1 | 109.1 | 172.9 | 0.0000 | 0.3255 | 0.1102 |
| Prostate cancer | Cancer | 30 | 30 | 0 | 100 | 67.2 | 10.1 | 25.4 | 3.6 | 76.4 | 125.8 | 176.3 | 0.0000 | 0.3790 | 0.0000 |
| Glomerulonephritis | Inflammation | 10 | 6 | 4 | 60 | 48.9 | 15.4 | 24.5 | 5.9 | 77.4 | 112.4 | 179.4 | 0.4914 | 0.8882 | 0.7209 |
| Cholecystitis | Inflammation | 30 | 17 | 13 | 56.7 | 65.9 | 10.3 | 24 | 3.6 | 76.3 | 145.1 | 159 | 0.0000 | 0.6204 | 0.7961 |
| Pancreatitis | Inflammation | 30 | 19 | 11 | 63.3 | 55.2 | 15.2 | 24.7 | 6.2 | 77.7 | 133.4 | 162.5 | 0.3908 | 0.7845 | 0.4348 |
| Hyperlipidemia | Lifestyle | 30 | 23 | 7 | 76.7 | 65.1 | 11.3 | 24.2 | 2.7 | 73.6 | 109.5 | 130.5 | 0.0000 | 0.5493 | 0.0596 |
| Hypertension | Lifestyle | 30 | 18 | 12 | 60 | 68.6 | 11.8 | 28.5 | 11.3 | 77.3 | 131.4 | 161.5 | 0.0000 | 0.0701 | 0.6042 |
| Diabetes type 1 | Lifestyle | 30 | 15 | 15 | 50 | 38.3 | 17.5 | 23.1 | 5.1 | 79.1 | 196.1 | 179.4 | 0.0001 | 0.2580 | 1.0000 |
| Diabetes type 2 | Lifestyle | 30 | 19 | 11 | 63.3 | 57.3 | 13.4 | 26.1 | 4.5 | 75.3 | 181.8 | 166.4 | 0.0770 | 0.1858 | 0.4348 |
| Obesity | Lifestyle | 30 | 22 | 8 | 73.3 | 60.3 | 16.2 | 29.7 | 2.7 | 74.3 | 121.5 | 153.8 | 0.0017 | 0.0000 | 0.1102 |

NC: Normal control

### Chemicals and reagents

HPLC-grade acetonitrile (ACN), water, and formic acid (FA) were purchased from Fisher Scientific (Loughborough, UK). Ammonium bicarbonate (ABC) was purchased from iNtRON Biotechnology (Seongnam, Korea). Dithiothreitol (DTT) was obtained from Amresco (Solon, OH, USA), iodoacetamide (IAA) from Sigma-Aldrich (St. Louis, MO, USA), and RapiGest SF surfactant from Waters Corporation (Milford, MA, USA). Sequencing-grade trypsin was purchased from Promega (Madison, WI, USA). For each endogenous (light) target peptide a corresponding stable-isotope-labelled (heavy) internal-standard peptide was synthesized with heavy-labelled residues giving a defined mass shift relative to the endogenous peptide. Heavy peptides were purified by HPLC, quantified by amino-acid analysis, and confirmed to be ≥ 98% pure by capillary zone electrophoresis; peptide peak areas were therefore not corrected for purity. Peptides were solubilized according to the manufacturer’s guidelines and verified to have the expected monoisotopic masses.

### Plasma sample preparation and digestion

A 2-µL volume of plasma was denatured and reduced by adding 0.2% RapiGest SF surfactant and 20 mM DTT in 100 mM ABC (pH 8.0); 20 µL of this stock solution was added to the sample, vortexed, and incubated at 60 °C for 60 min. The sample was then alkylated with 10 µL of 100 mM IAA, vortexed, and incubated in the dark at room temperature for 30 min. Digestion was initiated by adding 40 µL of trypsin solution (0.1 µg/µL), and the sample was vortexed and incubated at 37 °C for 4 h. Digestion was quenched by adding 10 µL of 10% FA, and insoluble material and surfactant by-products were removed by centrifugation at 15,000 rpm at 4 °C for 60 min. The supernatant was transferred to a fresh tube. All sample-preparation steps were performed on a Thermomixer C (Eppendorf, Westbury, NY, USA); total preparation time was approximately 12 h. Digested samples were transferred to an autosampler for MRM-MS analysis.

### Targeted MRM mass spectrometry

Chromatographic separation was performed on a fully automated online 1290 Infinity II liquid-chromatography system (Agilent Technologies, Santa Clara, CA, USA). The autosampler compartment was held at 4 °C and the LC separation at 40 °C. Sample clean-up used a guard column (2.1 × 30.0 mm, 1.8 µm, 80 Å) and peptides were separated on an analytical column (2.1 × 150.0 mm, 1.8 µm, 80 Å) (both Agilent Technologies), operated in parallel to increase throughput with mobile phases A (0.1% FA in water) and B (0.1% FA in ACN). Ten microlitres of digested sample was injected onto the guard column with the effluent to waste at 400 µL/min for 1 min in 10% B; after valve switching, flow was directed from the guard to the analytical column and 10% B was run at 40 µL/min for 1 min. Bound peptides were eluted on a linear gradient from 10% to 60% B over 5 min at 400 µL/min, followed by a column wash at 90% B for 1 min at 40 µL/min and re-equilibration at 10% B for 4 min. The total injection-to-injection cycle, including clean-up and re-equilibration, was 12 min, and the injector needle and tubing were washed after each injection with 50% aqueous ACN. Quantitative analysis was performed on an Agilent 6495 triple-quadrupole mass spectrometer with a Jet Stream electrospray source (Agilent Technologies), operated in positive-ion multiple-reaction-monitoring (MRM) mode. Source parameters were: gas temperature 250 °C, gas flow 15 L/min, nebulizer 30 psi, sheath-gas temperature 350 °C, and sheath-gas flow 12 L/min. The delta electron-multiplier voltage was 200 V, and the cell-accelerator and fragmentor voltages were 5 V and 380 V, respectively. The dwell and cycle times were 6 ms and 2,502 ms, respectively, and both Q1 and Q3 were set to unit resolution (0.7 Da at half height). The assay monitored 1,020 transitions covering 510 peptides from 126 target proteins selected for existing clinical or LDT relevance, each endogenous peptide paired with its heavy internal standard. Protein abundance was quantified as the peak-area ratio (PAR) of the endogenous light peptide to its heavy internal standard; provided PAR values equaled light area divided by heavy area to within numerical precision (maximum absolute difference 5 × 10 □ □ across all transitions), and PAR values were normalized across the 510 heavy peptides to control for sample loading.

### Quality control and matrix construction

Raw transition-level data comprised 250,390 quantified records across 490 samples, 126 proteins, 510 peptides, and 1,020 transitions (510 of which were the distinct light-transition quantitation keys used for matrix construction). Missing PAR values (0.39% overall) traced almost entirely to a single peptide, VCPFAGILENGAVR (apolipoprotein H, UniProt P02749), whose two transitions lacked a usable heavy internal standard and were undefined in all samples; this peptide was dropped and apolipoprotein H retained through its four other quantifiable peptides, leaving 509 usable peptides mapping to all 126 proteins with no missing values. Peptide-level abundance was computed as the median PAR across a peptide’s transitions and protein-level abundance as the median across a protein’s peptides (median three peptides per protein). Fourteen zero PAR values (endogenous peptide below the limit of detection; 10 peptides affected) were floored to half the minimum non-zero PAR of the corresponding column before log-transformation. Analytical reproducibility was assessed from the coefficient of variation of the heavy internal-standard peak areas across samples (median 41.5%, interquartile range 37.6–45.1%), consistent with crude-plasma targeted assays. Multivariate-outlier screening (PCA-distance z-score) flagged no sample at z > 5 and a single sample at z > 4, indicating a clean cohort (**Fig. S2**). Peaks were inspected in Skyline, with light and heavy peptides checked for co-elution of all transitions, retention-time alignment, relative-intensity correlation with the spectral library (dotp) and between light and heavy transitions (rdotp), and reproducibility across replicates.

### Normalization

Protein and peptide matrices were log □-transformed and centered by subtracting each sample’s median log □ PAR (per-sample median normalization). This reduced the between-sample spread of per-sample medians from a standard deviation of 0.356 to 0 while preserving biological differences (**Fig. S3**). Run-order drift was tested within each group by Spearman correlation of per-sample median abundance against replicate index; four of 17 groups showed nominal p < 0.05, none surviving multiple-testing correction.

### Unsupervised analysis

Principal-component analysis was performed on the z-scored 126-protein matrix (20 components retained). UMAP embedding used n_neighbors = 25, min_dist = 0.3, and a fixed random seed (42). Associations of the leading principal components with disease and super-class were tested by one-way ANOVA, and with age and sex by Pearson correlation. A group-mean abundance heatmap was clustered by Ward linkage (**Fig. 2**).

### Differential abundance

For each of the 16 diseases versus the normal-control group, protein-level differential abundance was assessed two ways: an unadjusted Mann–Whitney U test, and an age- and sex-adjusted ordinary-least-squares model (abundance ∼ disease indicator + age + sex, with median-imputed age and binary sex). p-values were corrected within each disease by the Benjamini–Hochberg procedure, and log □ fold-changes were computed relative to the control-group mean. Of 2,016 protein–disease tests, 607 were significant after age/sex adjustment (FDR < 0.05); adjustment retained the large majority of unadjusted signals, indicating that the associations are not demographic artefacts. Volcano plots for representative diseases are in **Fig. S4** and the full table is **Table S2**.

### Disease-detection classifiers

Per-disease detection used two model families under identical five-fold stratified cross-validation with out-of-fold prediction: an L2-regularised logistic-regression pipeline (standardization followed by logistic regression, max_iter = 2000, C = 1.0, balanced class weights), and gradient-boosted trees (XGBoost; 300 estimators, max_depth = 3, learning rate 0.05, subsample 0.8, colsample_bytree 0.6). Each disease was classified against the normal-control group, and performance was summarized by AUROC, area under the precision–recall curve, and sensitivity at 95% specificity. An any-disease-versus-healthy classifier pooled all disease samples against the 30 controls. Statistical significance of detection was assessed by permutation testing (200 label permutations per disease); all diseases yielded p ≤ 0.01 with permuted-null AUROC near 0.5. Per-disease ROC curves are in **Fig. S5** and performance values in **Table S3**.

### Multi-class disease identification

A single 17-class multinomial classifier (standardization followed by multinomial logistic regression, max_iter = 3000, C = 1.0, balanced class weights) assigned each sample to one of the 16 diseases or the control group under five-fold stratified cross-validation. Performance was summarized by balanced accuracy, macro-F1, and per-class precision, recall, and F1 (**Fig. 5, Fig. S6**). A gradient-boosted-tree alternative was evaluated for comparison (balanced accuracy 0.561 versus 0.584 for logistic regression), and the logistic-regression model was retained. Chance-level balanced accuracy is 1/17 = 0.059.

### Minimal panel and feature importance

Feature importance for the any-disease screen was computed as the mean absolute SHAP value from a gradient-boosted-tree model fitted to the full data. To construct the panel-size curve, proteins were ranked by SHAP importance and panels of increasing size (1–126 proteins) were re-evaluated under the same five-fold cross-validation for both the any-disease screen and the mean per-disease screen; per-disease SHAP-ranked top-five markers were also computed. The 20-protein consensus panel and per-disease markers are in **Table 2** and **Table S4**. Co-abundance structure was summarized as the median absolute inter-protein Pearson correlation (**Fig. S9**).

**Table 2.** Minimal 20-protein biomarker panel for multi-disease screening.

| Rank | Gene | UniProt | Mean_absSHAP | N diseases sig | Top5 marker |
| --- | --- | --- | --- | --- | --- |
| 1 | CXCL7 | P02775 | 0.5750 | 11 | Breast cancer; Colon cancer; Thyroid cancer; Pancreatic cancer; Bladder cancer; Kidney cancer; Prostate cancer; Glomerulonephritis; Cholecystitis; Pancreatitis; Obesity |
| 2 | ANGT | P01019 | 0.5650 | 3 | Diabetes type 1; Diabetes type 2 |
| 3 | MET | P08581 | 0.5000 | 9 | Glomerulonephritis; Hyperlipidemia |
| 4 | PLF4 | P02776 | 0.4680 | 11 | Breast cancer; Colon cancer; Thyroid cancer; Pancreatic cancer; Bladder cancer; Kidney cancer; Prostate cancer; Glomerulonephritis; Cholecystitis; Pancreatitis; Obesity |
| 5 | ITA2B | P08514 | 0.4300 | 11 | Colon cancer; Thyroid cancer; Bladder cancer; Kidney cancer; Prostate cancer; Cholecystitis; Obesity |
| 6 | RENI | P00797 | 0.4010 | 13 | Kidney cancer; Prostate cancer; Hypertension; Obesity |
| 7 | VINC | P18206 | 0.3800 | 9 | Kidney cancer; Prostate cancer; Cholecystitis; Hyperlipidemia |
| 8 | CD14 | P08571 | 0.3600 | 12 | Pancreatic cancer; Bladder cancer; Pancreatitis; Hypertension |
| 9 | KIT | P10721 | 0.2790 | 1 | Hyperlipidemia |
| 10 | FINC | P02751 | 0.2440 | 9 | Breast cancer; Colon cancer; Thyroid cancer; Hypertension; Diabetes type 2 |
| 11 | BTD | P43251 | 0.2250 | 6 | Hypertension |
| 12 | ICAM1 | P05362 | 0.2010 | 3 | — |
| 13 | ALBU | P02768 | 0.1800 | 11 | Pancreatic cancer; Glomerulonephritis; Pancreatitis; Diabetes type 1 |
| 14 | MSH2 | P43246 | 0.1750 | 7 | Hypertension; Diabetes type 1; Diabetes type 2 |
| 15 | CO7 | P10643 | 0.1610 | 5 | Liver cancer |
| 16 | F13A | P00488 | 0.1590 | 8 | — |
| 17 | FETA | P02771 | 0.1510 | 5 | — |
| 18 | LDHB | P07195 | 0.1250 | 8 | Colon cancer; Thyroid cancer; Cholecystitis |
| 19 | AMBP | P02760 | 0.1220 | 3 | — |
| 20 | A1AG1 | P02763 | 0.1180 | 3 | Hyperlipidemia |

### Robustness analyses

Two robustness analyses address the interpretation of the high detection performance. First, to test dependence on demographics, each protein was residualized on age and sex (linear regression) and the per-disease classifiers were re-run on the residuals; mean AUROC changed by only 0.003 (**Fig. S8**), indicating that the signal is not carried by age or sex. Second, to test whether high performance is specific to the proteome, we substituted routine clinical-chemistry and haematology values as classifier features; these also discriminated most diseases at near-ceiling AUROC (**Fig. S7**), which we interpret as evidence of case-control batch structure that any sufficiently rich feature set can exploit. A full account of the interpretation of near-perfect within-cohort discrimination is given in the Supplementary Text.

### Software

Analyses were performed in Python with scikit-learn (1.9.0), statsmodels (0.14.6), XGBoost (3.3.0), UMAP-learn (0.5.12), SHAP (0.52.0), SciPy (1.18.0), pandas (3.0.3), and NumPy (2.4.6). Mass-spectrometry peaks were inspected in Skyline. A fixed random seed (42) was used throughout.

## Results

### A targeted MRM assay quantifies 126 plasma proteins across 16 diseases

The overall study design—from cohort and single-plasma sampling through targeted MRM-MS quantitation to the integrated analyses and their outcome—is summarized in **Fig. 1**. In brief, one plasma sample per participant is measured on a single targeted assay, and the resulting quantitative proteome feeds five analyses (unsupervised structure, differential abundance, per-disease screening, disease identification, and minimal-panel selection) that together establish the feasibility of multiplexed proteomic screening.

**Fig. 1.**
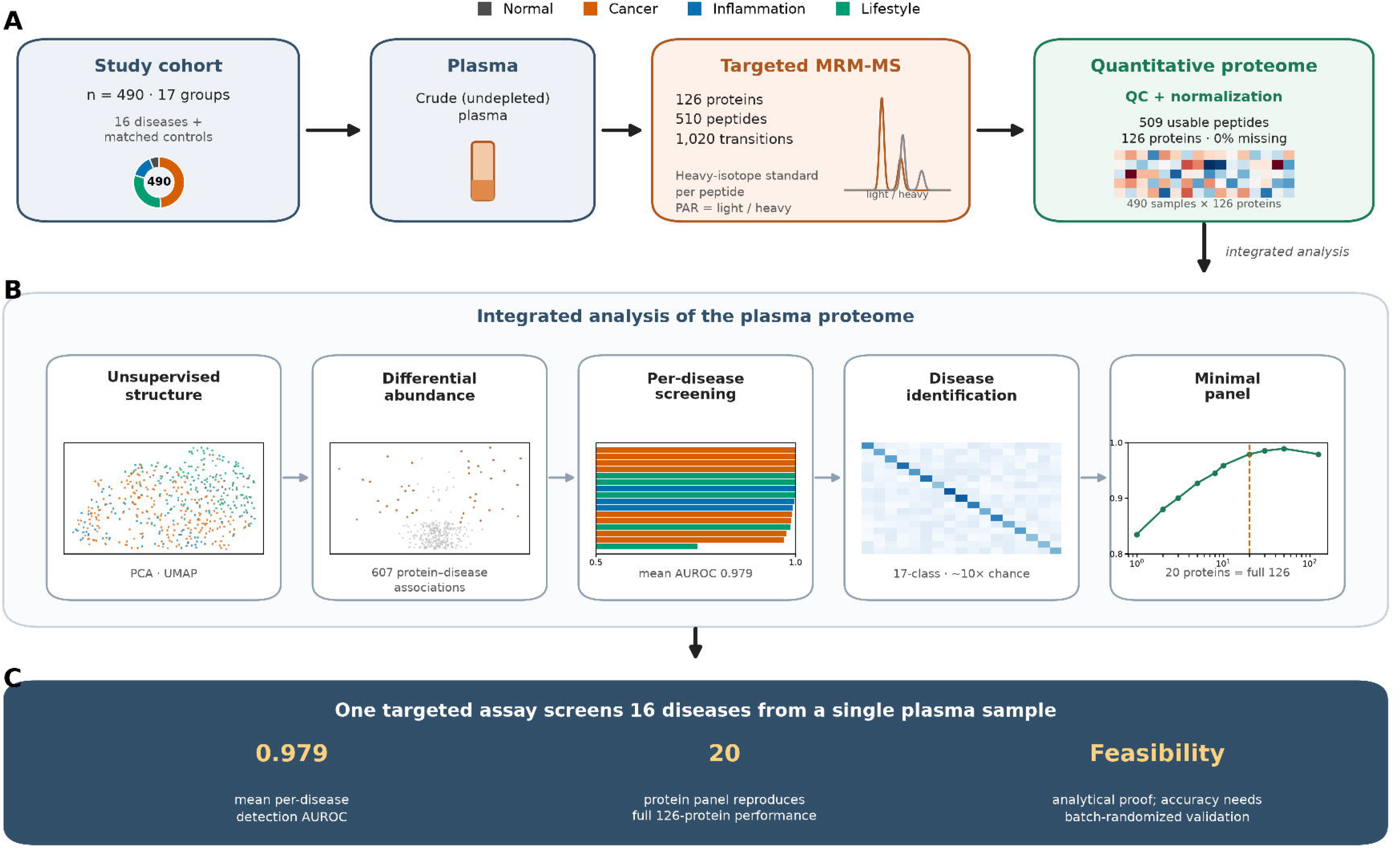
Study workflow: one targeted assay for simultaneous multi-disease screening. A graphical overview of the study, read top to bottom. (A) Experimental pipeline: a cohort of 490 participants in 17 groups (donut, colored by super-class; 16 diseases plus matched normal controls) each contributes a single crude-plasma sample, analyzed by targeted MRM-MS (126 proteins, 510 peptides, 1,020 transitions; each peptide quantified against a stable-isotope heavy internal standard as the peak-area ratio, PAR = light/heavy area) to yield, after QC and normalization, a complete quantitative proteome matrix (509 usable peptides, 126 proteins, no missing values). (B) Integrated analysis of the proteome through five modules—unsupervised structure (PCA/UMAP), differential abundance versus matched controls, per-disease screening classifiers, 17-class disease identification, and minimal-panel selection—with a representative result inset for each. (C) Principal outcome: a single assay detects the 16 diseases at a mean cross-validated AUROC of 0.979, a 20-protein subset reproduces the full 126-protein performance, and the results establish analytical feasibility while requiring batch-randomized prospective validation for clinical-accuracy claims. Colors denote super-class throughout (gray, normal; orange, cancer; blue, inflammation/renal; green, lifestyle/metabolic).

We assembled a cohort of 490 participants in 17 groups: a normal-control group (n = 30) and 16 disease groups (n = 30 each, except glomerulonephritis n = 10), organized into four super-classes— cancer (eight types, 240 participants), lifestyle or metabolic disease (five types, 150), inflammatory and renal disease (three types, 70), and normal controls (30). The normal-control group was demographically representative of the disease groups (age 52.4 ± 6.6 years, 15 men and 15 women, body-mass index 24.2 kg/m²), providing a matched comparison rather than a young-healthy-volunteer baseline (**Table 1; Fig. S1**).

Each plasma sample was analyzed by targeted MRM-MS monitoring 1,020 transitions covering 510 peptides from 126 proteins, with a stable-isotope-labelled heavy peptide for every target. Protein abundance was quantified as the peak-area ratio (PAR) of the endogenous (light) peptide to its heavy internal standard, normalized across the 510 heavy peptides. This internal-standard-referenced design yields a directly comparable, absolute abundance for each protein in each sample. The raw acquisition comprised 250,390 quantified transition-level records across the cohort.

After quality control (**Fig. S2**), one peptide (VCPFAGILENGAVR, apolipoprotein H) lacked a usable heavy standard and was removed, leaving 509 quantified peptides mapping to all 126 proteins with no missing values (overall missingness before removal, 0.39%). Peptide-level abundance was summarized as the median across a peptide’s transitions and protein-level abundance as the median across a protein’s peptides (median three peptides per protein). The median coefficient of variation of the heavy internal standards across samples was 41.5% (interquartile range 37.6–45.1%), consistent with crude-plasma targeted assays, and multivariate-outlier screening flagged no sample beyond a conservative threshold (**Fig. S2**). Log-transformation and per-sample median normalization removed a modest per-sample loading offset—reducing the between-sample spread of per-sample medians essentially to zero—without introducing run-order drift (**Fig. S3**). Together these results establish a reproducible, quantitative multi-disease proteome resource on which the analyses below are built.

### Plasma proteome variation is organized by disease rather than demographics

To ask what structures this proteome, we performed principal-component analysis (PCA) and uniform-manifold-approximation projection (UMAP) on the normalized 126-protein matrix (**Fig. 2A, B**). Samples separated by disease and by super-class, and a group-mean abundance heatmap resolved coherent, disease-specific protein signatures (**Fig. 2C**). The leading principal components were strongly associated with disease (one-way ANOVA F = 18.0 for PC1, p = 1.7 × 10 ³) and with super-class, but were essentially uncorrelated with age or sex (|r| ≤ 0.16 for the top components). Age and sex therefore contribute little to the dominant axes of proteome variation, which are instead disease-driven. This separation is a prerequisite for using the proteome as a disease screen rather than as a demographic readout, and it indicates that the structure the classifiers below exploit is not merely a reflection of the age or sex composition of the groups.

**Fig. 2.**
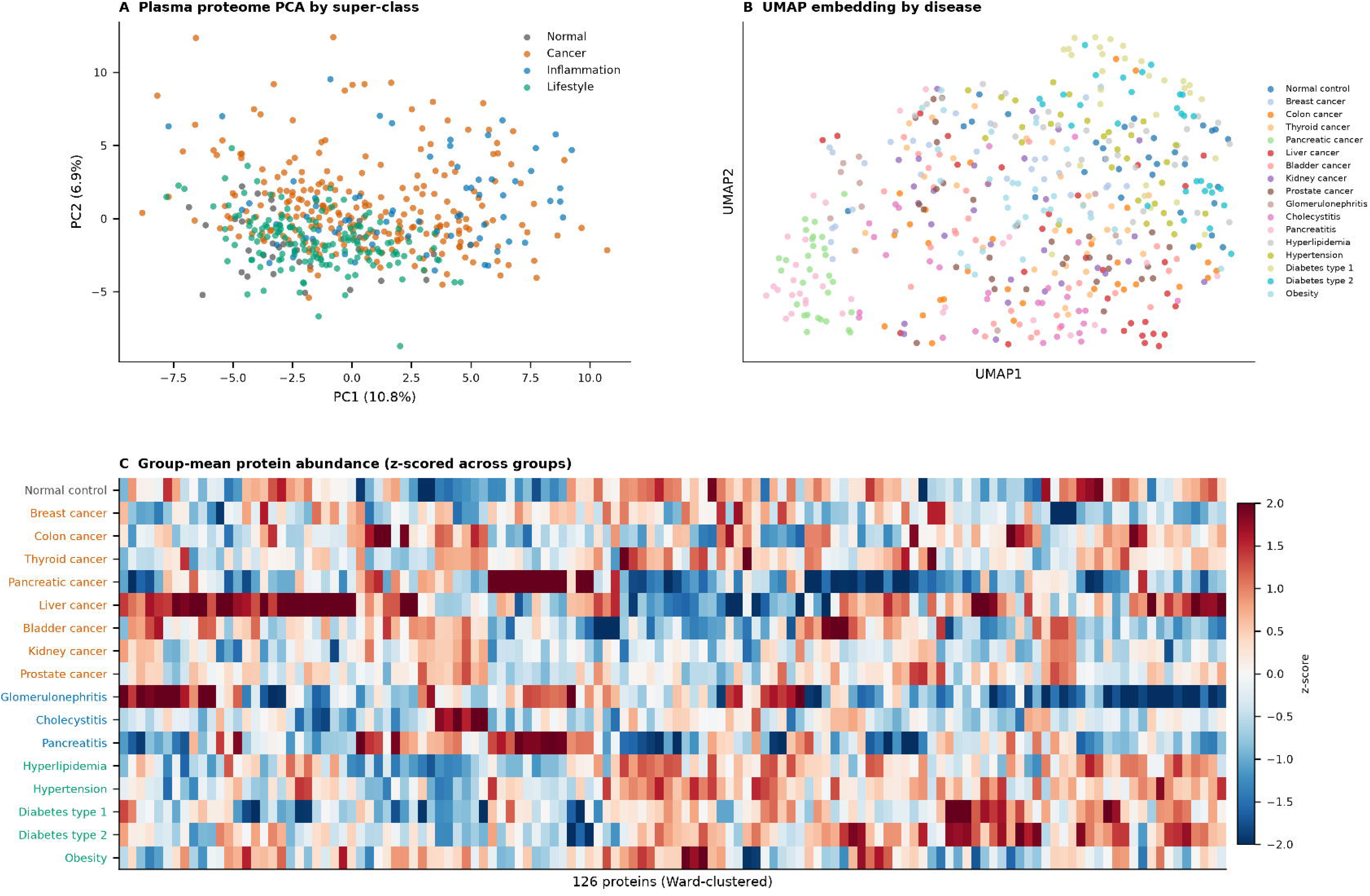
Plasma proteome variation is organised by disease rather than demographics. (A) Principal-component analysis (PCA) of the normalised 126-protein matrix (n = 490 samples), coloured by super-class; the first two components explain 10.8% and 6.9% of the total variance. Cancer and inflammatory/renal samples spread away from the tightly clustered normal and metabolic/lifestyle samples along PC1. (B) UMAP embedding of the same matrix (n_neighbors = 25, min_dist = 0.3, random seed = 42), coloured by individual disease; several conditions—most visibly pancreatic cancer—form coherent regions, while related diseases overlap. (C) Ward-clustered heatmap of group-mean protein abundance, z-scored across the 17 groups (rows) for all 126 proteins (columns); red and blue denote group-mean abundance above and below the cross-group average. Disease-specific blocks of concordantly high or low proteins are evident (for example the liver-cancer and glomerulonephritis rows), showing that the dominant structure in the proteome is disease-related. The leading components were strongly associated with disease (one-way ANOVA F = 18.0 for PC1, p = 1.7 × 10 □³ □) but essentially uncorrelated with age or sex (|r| ≤ 0.16 for the top components).

### Individual diseases are detected against matched controls with high accuracy

We next trained classifiers to distinguish each disease from the normal-control group, using L2-regularised logistic regression on the 126-protein matrix with five-fold stratified cross-validation and out-of-fold prediction; gradient-boosted trees were evaluated in parallel as an independent model family (**Figs. 3, 4**). Differential-abundance analysis (Mann–Whitney U tests together with age- and sex-adjusted linear models, Benjamini–Hochberg correction) identified 607 significant protein– disease associations after adjustment (**Fig. 3A, B; Fig. S4**), including disease-specific markers such as platelet factor 4 (PLF4) in pancreatic cancer, CXCL7 in pancreatitis, and albumin in glomerulonephritis (**Fig. 3C**). Age and sex adjustment retained the large majority of unadjusted signals, indicating that the associations are not demographic artefacts. Detection performance was high across the board: the mean cross-validated AUROC was 0.979, the median sensitivity at 95% specificity was 1.0, and 7 of 16 diseases reached AUROC = 1.000 (**Fig. 4B, C; Fig. S5**). Hyperlipidaemia was the weakest (AUROC 0.754), consistent with a proteome signature that overlaps substantially with matched controls. A single any-disease-versus-healthy classifier, pooling all disease samples against the 30 controls, achieved AUROC 0.973 (**Fig. 4A**). Permutation testing (200 label permutations per disease) confirmed that every disease was detected far above chance (all p ≤ 0.01; permuted-null AUROC ≈ 0.5; **Fig. 4D**). These results demonstrate that, within this cohort, the multiplexed proteome carries sufficient information to flag the presence of each individual disease against a matched control group.

**Fig. 3.**
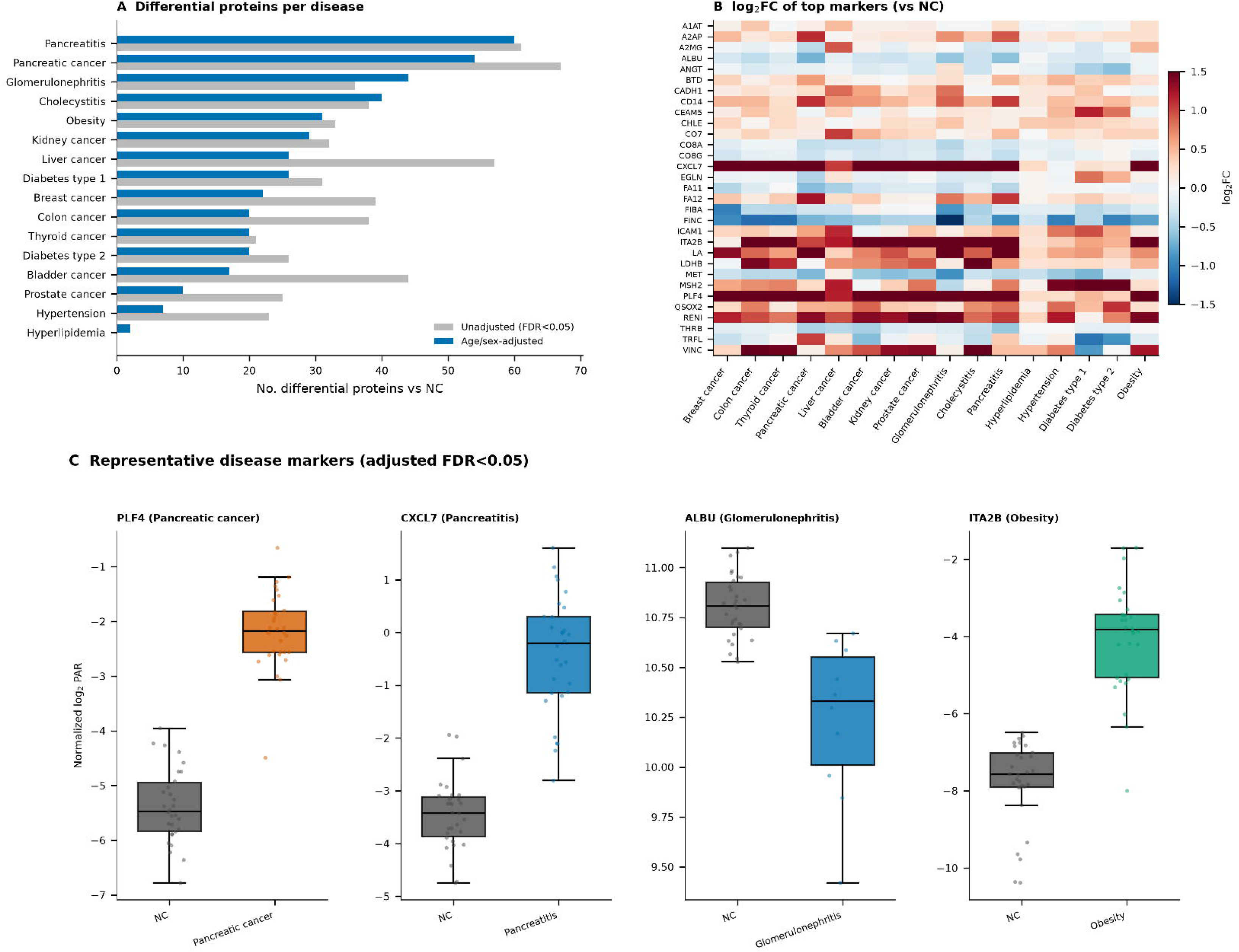
Differential protein abundance across diseases. (A) Number of proteins significantly different from matched normal controls for each disease, comparing an unadjusted Mann–Whitney U test (grey) with an age- and sex-adjusted linear model (blue, abundance ∼ disease + age + sex); adjustment retains the large majority of signals, indicating that the associations are not demographic artefacts. Diseases are ordered by adjusted count. (B) Heatmap of log □ fold-change (versus normal control) for a curated set of top markers (rows) across all 16 diseases (columns); red denotes higher and blue lower abundance in disease. Broadly elevated markers such as CXCL7, PLF4, ITA2B, and RENI appear as near-continuous red rows, whereas FINC and FIBA are broadly reduced. (C) Distributions of four representative, disease-specific markers in normal controls (NC) versus the relevant disease group, each significant after age/sex adjustment (FDR < 0.05): PLF4 in pancreatic cancer, CXCL7 in pancreatitis, albumin (ALBU) in glomerulonephritis, and ITA2B in obesity. Boxes show the median and interquartile range, whiskers extend to 1.5× IQR, and points are individual participants (normalised log □ PAR). Across all diseases, 607 of 2,016 protein–disease comparisons were significant after adjustment (Benjamini–Hochberg FDR < 0.05).

**Fig. 4.**
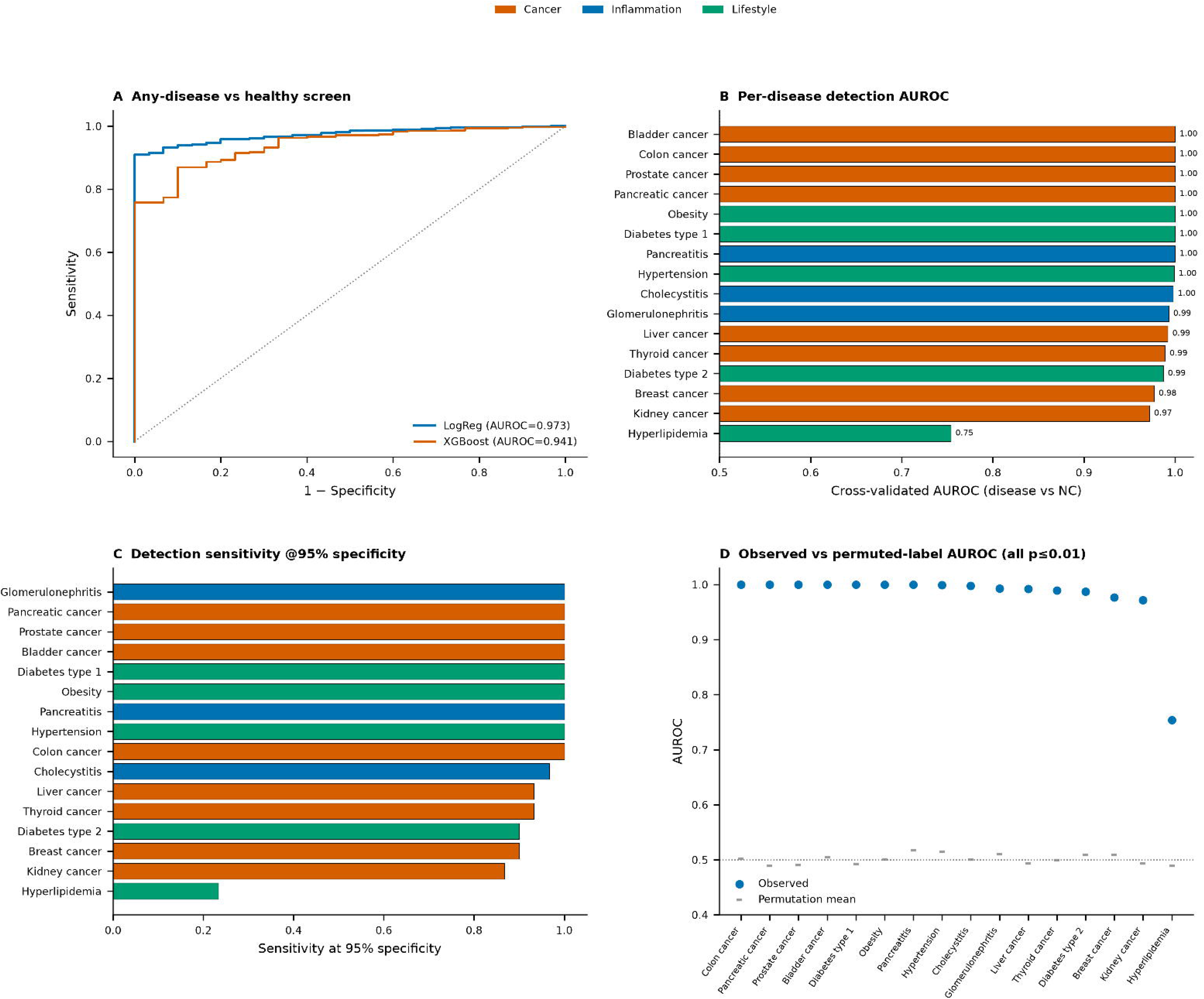
Per-disease detection performance. (A) Receiver-operating-characteristic (ROC) curves for the single any-disease-versus-healthy classifier, pooling all disease samples against the 30 normal controls, for L2-regularised logistic regression (AUROC = 0.973) and gradient-boosted trees (AUROC = 0.941); the diagonal marks chance. (B) Cross-validated per-disease detection AUROC (each disease versus matched controls; five-fold stratified cross-validation with out-of-fold prediction), coloured by super-class and annotated with values; 7 of 16 diseases reach AUROC = 1.00 and hyperlipidaemia is the weakest at 0.75. (C) Sensitivity at a fixed 95% specificity for each disease, a threshold-based summary more relevant to screening than AUROC alone; the median across diseases is 1.0. (D) Observed AUROC (filled points) versus the mean of 200 label permutations (dashes) for each disease; every disease exceeds its permuted null (all p ≤ 0.01), with permuted-null AUROC clustered near 0.5. Because permutation preserves the case-control batch structure, it tests only against chance label association and not against technical confounding.

### A single multinomial model identifies the specific disease

Beyond detecting that some disease is present, we asked whether the specific disease could be named. A single 17-way multinomial classifier assigned each sample to one of the 16 diseases or the control group with a balanced accuracy of 0.584 and a macro-F1 of 0.589—about ten times the chance level of 0.059 (**Fig. 5**). Discrimination was strong for cancers with distinct proteome signatures (liver cancer F1 = 0.81, pancreatic cancer 0.81) and for metabolic disease (obesity 0.78, type-1 diabetes 0.72), and weaker where signatures overlapped: kidney and prostate cancer were most often confused with each other, and the two diabetes subtypes with each other (**Fig. 5; Fig. S6**). Crucially, this confusion structure is biologically interpretable rather than random, indicating that the residual errors reflect genuine proteomic similarity between related conditions rather than a failure of the classifier. A gradient-boosted-tree alternative gave comparable but slightly lower balanced accuracy (0.561 versus 0.584), and the logistic-regression model was retained.

**Fig. 5.**
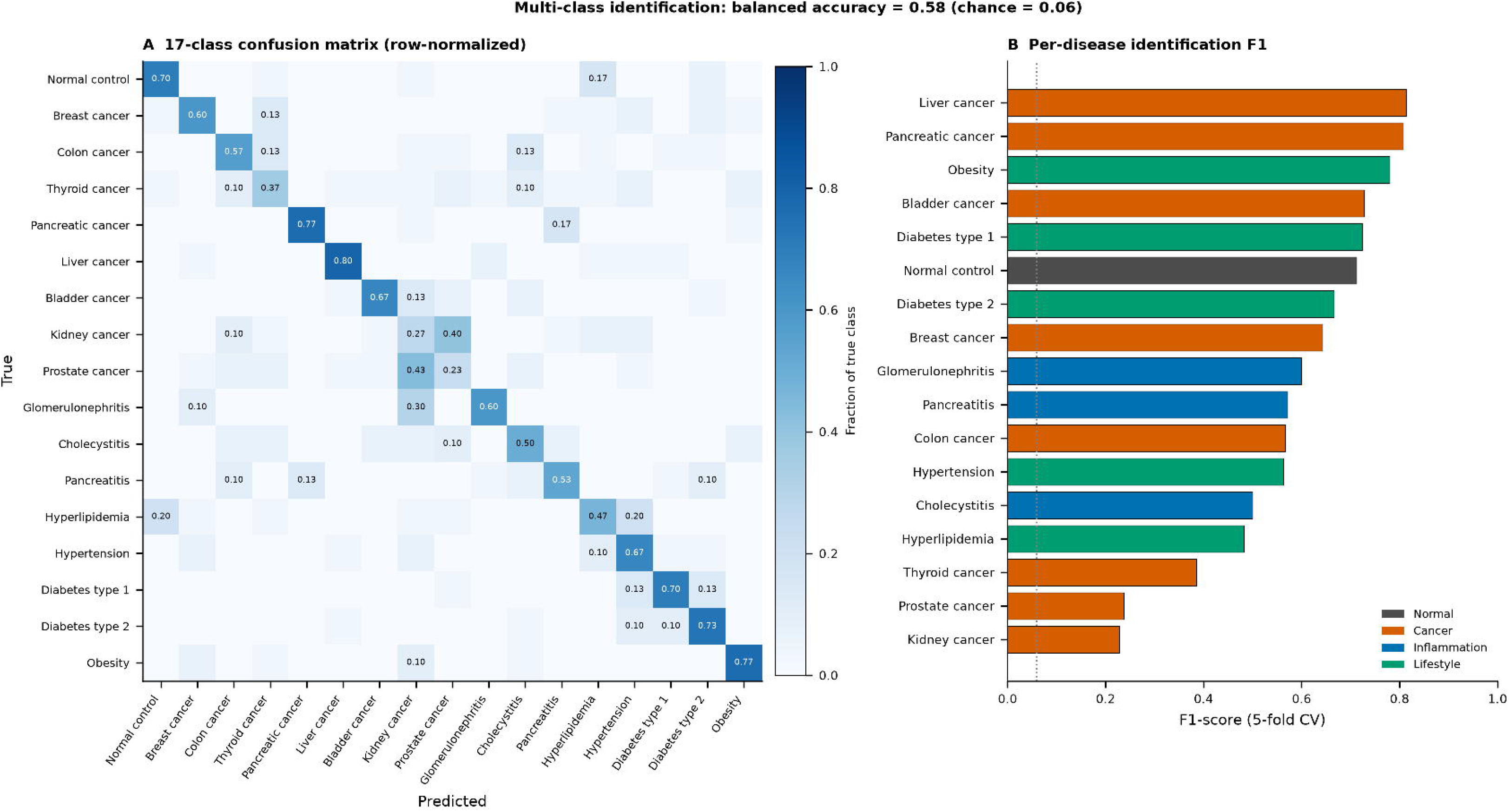
Specific disease identification. (A) Row-normalised confusion matrix for a single 17-class multinomial classifier assigning each sample to one of the 16 diseases or the normal-control group (five-fold stratified cross-validation); cell values are the fraction of each true class (rows) predicted as each class (columns), so the diagonal gives per-class recall. Overall balanced accuracy is 0.58 against a chance level of 0.06 (1/17). Off-diagonal mass is concentrated between biologically related conditions—kidney versus prostate cancer, and the two diabetes subtypes—rather than scattered at random, indicating that residual errors reflect genuine proteomic similarity. (B) Per-disease F1 score (harmonic mean of precision and recall) from the same model, coloured by super-class and ordered by value; identification is strongest for cancers and metabolic diseases with distinctive proteomes (liver and pancreatic cancer, obesity, type-1 diabetes) and weakest where signatures overlap (kidney and prostate cancer).

### A minimal 20-protein panel reproduces full-panel performance

For a screen to be practical, the protein list should be as short as possible. Ranking proteins by SHAP importance in the any-disease model and re-evaluating panels of increasing size revealed sharply diminishing returns (**Fig. 6**): a 20-protein subset reached a mean per-disease AUROC of 0.979— matching the full 126-protein panel—and even 10 proteins reached 0.959. The consensus panel was led by platelet-derived and vascular proteins (CXCL7, PLF4), the renin–angiotensin components angiotensinogen (ANGT) and renin (RENI), hepatocyte growth factor receptor (MET), and integrin and adhesion proteins (ITA2B, VINC, ICAM1) (**Table 2**). Co-abundance analysis showed that the panel is largely non-redundant (median inter-protein |r| = 0.085; Fig. S9), so each protein contributes largely independent information. A 20-protein assay of this kind would be substantially cheaper and faster to deploy than the full panel while preserving its discriminative content.

**Fig. 6.**
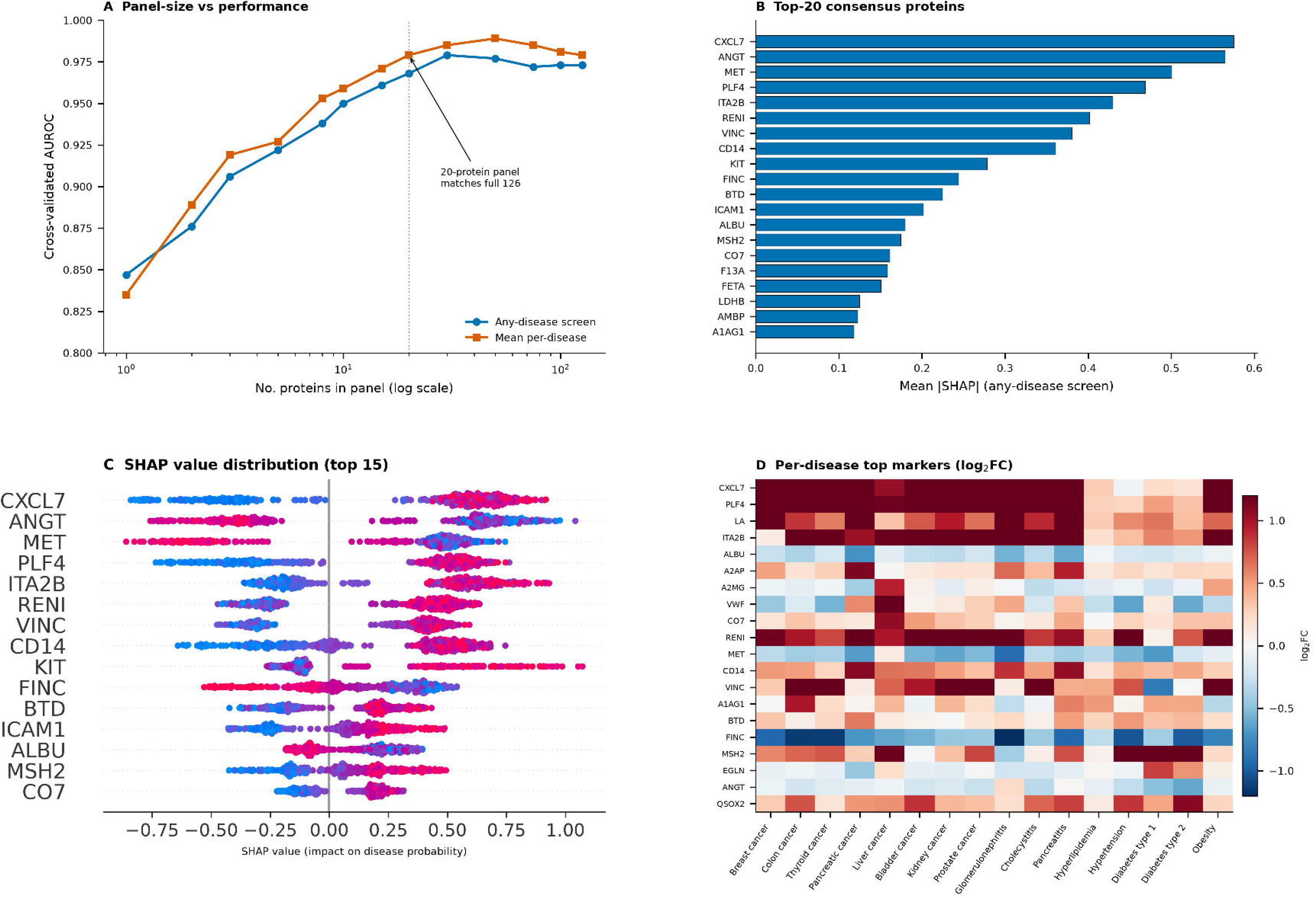
A minimal 20-protein panel reproduces full-panel performance. (A) Cross-validated AUROC as a function of panel size, for the any-disease screen (blue) and the mean per-disease screen (orange), with proteins added in order of decreasing SHAP importance (x-axis log-scaled). Performance rises steeply then plateaus: a 20-protein panel (dotted line) matches the full 126-protein panel, and 10 proteins already reach AUROC ≈ 0.96. (B) The top-20 consensus proteins ranked by mean absolute SHAP value in the any-disease screen, led by CXCL7, ANGT, MET, PLF4, and ITA2B. (C) SHAP-value distributions for the 15 most important proteins (each point a sample, coloured by that protein’s abundance), showing the direction in which high or low abundance drives the disease prediction. (D) Heatmap of per-disease log □ fold-change for the panel proteins across all 16 diseases, showing that a compact, largely non-redundant set (median inter-protein |r| = 0.085) carries disease-discriminating information spanning cancers, inflammatory/renal, and metabolic disease.

### Near-perfect discrimination represents an optimistic upper bound

The per-disease AUROCs reported above are within-cohort estimates, and several converging lines of evidence indicate that they are optimistic upper bounds rather than deployable accuracies. First, the problem is high-dimensional relative to sample size (126 features, roughly 60 samples per binary task), a regime in which cross-validated performance is upwardly biased because feature-rich models can exploit incidental structure that does not generalize. Second, and more important, each disease group was acquired as a separate case-control batch; any systematic analytical difference between batches is perfectly confounded with disease status, and label permutation—which preserves batch structure— cannot detect it. A permutation p ≤ 0.01 therefore confirms that the discrimination is not a chance artefact, but cannot establish that it is biological rather than technical. Third, when we substituted routine clinical-chemistry values (glucose, lipids, liver enzymes, and standard haematology) for the proteome, these too discriminated most diseases at near-ceiling AUROC (**Fig. S7**)—an implausible result for genuine early screening that points to batch or selection structure any sufficiently rich feature set can exploit. The proteome signal is nonetheless real in the biological sense. Performance was essentially unchanged after removing age and sex variance by residualization (mean AUROC drop 0.003; **Fig. S8**), so it is not a demographic artefact; and the marker biology is coherent and disease-specific (platelet factor 4 and CXCL7 as broad markers of malignancy and inflammation; angiotensinogen and renin in hypertension and diabetes; albumin in glomerulonephritis), consistent with known pathophysiology rather than arbitrary. The analytical platform—reproducible, internal-standard-referenced multiplexed quantitation of 126 proteins in crude plasma with no missing data—performs exactly as intended. We therefore interpret the analytical results as establishing feasibility (reproducible multiplexed quantitation and disease-structured signal) while explicitly deferring accuracy claims to prospective, multi-batch validation.

## Discussion

We have shown that a single targeted mass-spectrometry assay can quantify 126 plasma proteins reproducibly in crude plasma across a 490-participant, 16-disease cohort, and that the resulting proteome is organized by disease biology to a degree that supports both detection and specific identification of disease in cross-validation. A 20-protein subset captures essentially the full signal, which is encouraging for a low-cost, deployable assay. These are the analytical foundations that a multiplexed proteomic screen would require: reproducible quantitation across a wide abundance range, no missing data, coherent disease-specific biology, and a compact feature set. To our knowledge, this is among the broadest single-assay protein readouts assembled across cancer, inflammatory or renal, and metabolic disease simultaneously, rather than within a single disease category or a single super-class^4^.

The central caveat is validation. Our cohort was assembled as disease-specific case-control batches without an independent, prospectively collected validation set, and we have shown that this structure alone can produce near-perfect within-cohort discrimination—including from routine laboratory values that could not plausibly screen these diseases de novo. This is not a peculiarity of our study but a general and well-documented hazard in biomarker research, and it is the reason that so few discovery-stage classifiers survive external testing^10, 11, 14^. Batch and pre-analytical structure that is perfectly aligned with the outcome cannot be removed by cross-validation, permutation testing, or demographic adjustment, because each of these operates within the existing sample set and therefore preserves the confound^38^. The honest reading of our data is therefore feasibility, not clinical accuracy: the assay is analytically sound and the biology is real, but the performance numbers must be re-established in a design that breaks the disease–batch confound.

Two findings are nevertheless robust to these concerns and worth emphasizing. First, the proteome signal is not a demographic readout: removing all age and sex variance leaves classification essentially unchanged, and the dominant axes of proteome variation are uncorrelated with age and sex. Second, the differential-abundance biology recapitulates known pathophysiology—platelet-derived and inflammatory chemokines in malignancy and pancreatitis, renin–angiotensin components in hypertension and diabetes, and albumin loss in glomerulonephritis—rather than arbitrary features. The analytical reproducibility we observe is consistent with the established performance of internal-standard-referenced MRM assays, which transfer between laboratories more reliably than untargeted discovery workflows^18, 28^.

Three steps follow directly and define the path to a clinically meaningful claim. First, prospective sample collection in which disease and control samples are randomized across acquisition batches, so that batch cannot proxy for diagnosis; without this, no within-cohort accuracy—however high—can be trusted. Second, an independent validation cohort, ideally drawn from a screening-relevant population (asymptomatic individuals at routine examination) rather than confirmed patients, because screening performance and diagnostic performance differ substantially and the former is what a population screen must deliver^11^. Third, benchmarking against—and integration with—the routine clinical chemistry that already accompanies a health check, to establish the proteome’s incremental value over inexpensive standard labs rather than its performance in isolation. Standardized frameworks for analytical validation of multiplex assays and for transparent reporting of prediction models provide a template for each of these steps^39, 40^.

The broader motivation is economic and logistical. If a compact, internal-standard-referenced protein panel can be measured in crude plasma on instrumentation already deployed in clinical laboratories, a single injection could in principle report on many organ systems at a marginal cost far below that of the dozens of separate assays it would replace^21, 36^, and could complement nucleic-acid multi-cancer early-detection tests that by construction do not address common non-malignant disease. The 20-protein panel identified here, dominated by non-redundant platelet, vascular, renin–angiotensin, and adhesion proteins, is a plausible starting point for such an assay. Realizing that potential, however, depends entirely on the validation design above; the contribution of the present work is to demonstrate that the measurement is reproducible and the signal is disease-structured, and to define— quantitatively and explicitly—the confound that any subsequent accuracy claim must overcome.

In summary, a single MRM plasma-proteome assay reproducibly quantifies a medically relevant, 126-protein multiplex in crude plasma and resolves disease-structured biological signal across sixteen diverse conditions, with a 20-protein subset preserving the discriminative content. The within-cohort accuracies are optimistic upper bounds set by case-control batch structure, and we bound this limitation directly. The reproducible measurement demonstrated here is the foundation on which a batch-randomized, independently validated multi-disease screen can now be built and tested as what it is designed to be: a single assay that screens for many diseases at once.

## Data and Code availability

The mass spectrometry data have been deposited to Panorama Public (https://panoramaweb.org/), and are accessible at https://panoramaweb.org/51tcZP.url. All analysis scripts used in this study, including cosine similarity calculation, differential abundance analysis (limma), and figure generation, are publicly available on GitHub (https://github.com/kimlab-cnu/MultiDiseaseMarker). Additional data are available from the corresponding author on reasonable request.

## Funding

This work was supported by the National Research Foundation of Korea (NRF) grants funded by the Korean government (MSIT) (RS-2023-00209456, RS-2025-24803258, and RS-2026-25488704), and by the Korea Basic Science Institute (National Research Facilities and Equipment Center) grant funded by the Korean government (MSIT) (RS-2024-00402298). This work was also supported by the Basic Science Research Program through the National Research Foundation of Korea (NRF) funded by the Ministry of Education (RS-2025-25436019), and by a grant from the Ministry of Food and Drug Safety (RS-2024-00331799).

## Acknowledgement

The biospecimens and data used for this study were provided by the Biobank of Chungbuk National University Hospotal (CBNUH), Jeonbuk National University Hospital (JBUH), Korea Atomic Energy Research Institute (KAERI), Asan Medical Center (AMC) and Inje University Busan Paik Hospital (BPH), a member of the Korea Biobank Network. All materials derived from the Biobank of Korea were obtained (with informed consent) under institutional review board (IRB)-approved protocols. We acknowledge the use of Claude Opus 4.8 solely for linguistic refinement and grammatical corrections in manuscript preparation. All scientific content, data analysis, and intellectual contributions presented herein were developed independently by the authors without the use of generative AI tools.

## Author Contributions

A.S. and H.K.: conceptualization, methodology, experimental analysis, data analysis, visualization; J.J., E.H., Y.C., J.P., H.L., and S.P.: data analysis; A.S, and H.K.: writing-original draft, conceptualization, project administration, resources, supervision, writing-review & editing. All authors have read and approved the final manuscript.

## Conflicts of Interest

The authors declare no conflicts of interest.

## Notes

### Competing Interest Statement

The authors have declared no competing interest.

### Summary of Updates

Distribution/Reuse options updated: All Rights Reserved: By selecting this option, you are retaining all rights over the use of your article. Anyone who wishes to share, reuse, remix, or adapt your article must first contact you for permission.

