## Supplemental Text for "A targeted mass-spectrometry plasma proteome for simultaneous multi-disease screening"

**Interpretation of near-perfect within-cohort discrimination**

The per-disease detection AUROCs reported in the main text (mean 0.979, seven diseases at 1.000) are exceptional and warrant careful interpretation, because effect sizes this large in a biomarker study are more often a signature of study-design confounding than of deployable diagnostic accuracy. We lay out the evidence and our reasoning explicitly.

***High dimensionality relative to sample size***

Each binary detection task compares roughly 30 disease samples with 30 controls using 126 protein features (509 at the peptide level). In this p ≫ n regime, even cross-validated performance is upwardly biased, because feature-rich models can exploit incidental structure that does not generalise. Nested cross-validation and permutation testing mitigate but do not eliminate this bias.

***Case-control batch structure***

The decisive concern is that each disease group was assembled and acquired as a distinct case-control batch. Mass-spectrometry measurements are sensitive to acquisition conditions (instrument state, calibration, sample age, processing date). When all samples of a given disease share a batch that differs from the control batch, any systematic analytical difference between batches is perfectly confounded with disease status. Critically, label-permutation testing does not detect this: permuting labels within the existing sample set preserves the batch structure, so the permuted null reflects only the absence of a random label association, not the absence of a batch association. A p ≤ 0.01 permutation result therefore confirms the discrimination is not a chance artefact, but cannot confirm it is biological rather than technical.

***The clinical-chemistry benchmark***

The most direct evidence that batch/selection structure is present comes from substituting routine clinical-chemistry and haematology values—glucose, lipids, liver enzymes, complete blood count—for the proteome. These standard laboratory values also discriminated most diseases at near-ceiling AUROC. It is not biologically plausible that a fasting glucose and lipid panel can distinguish, for example, bladder cancer from matched controls at AUROC 0.95 by genuine screening biology. The most parsimonious explanation is that the cohort carries case-control structure (batch, selection, or pre-analytical) that any sufficiently rich feature set can read out.

***What is nonetheless established***

Two results are robust to these concerns. First, the proteome signal is not a demographic artefact: removing all age and sex variance by residualisation leaves performance essentially unchanged (mean AUROC drop 0.003; **Fig. S8**), and the leading axes of proteome variation are uncorrelated with age and sex (**Fig. 2**). Second, the differential-abundance biology is coherent and disease-specific (platelet factor 4 and CXCL7 as broad markers of malignancy and inflammation; the renin–angiotensin components angiotensinogen and renin in hypertension and diabetes; albumin in glomerulonephritis), consistent with known pathophysiology rather than arbitrary. The analytical platform—reproducible internal-standard-referenced multiplexed quantitation of 126 proteins in crude plasma with no missing data—performs as intended.

***Conclusion***

We therefore frame the contribution as analytical feasibility, not clinical accuracy. The assay reproducibly quantifies a medically relevant multiplexed proteome, and that proteome carries real, disease-structured biological signal. Establishing screening accuracy requires a design that breaks the disease-batch confound: prospective collection with batch-randomised diseases and controls, and an independent validation cohort drawn from a screening-relevant (asymptomatic) population. These are the necessary next steps, and the reproducible measurement demonstrated here is their foundation.

**Underpowered glomerulonephritis group**

The glomerulonephritis group (n = 10) is underpowered relative to the other groups (n = 30). Its detection and identification metrics should be regarded as provisional, and it was retained primarily to preserve the breadth of the inflammatory/renal super-class.

**Supplementary Figures**

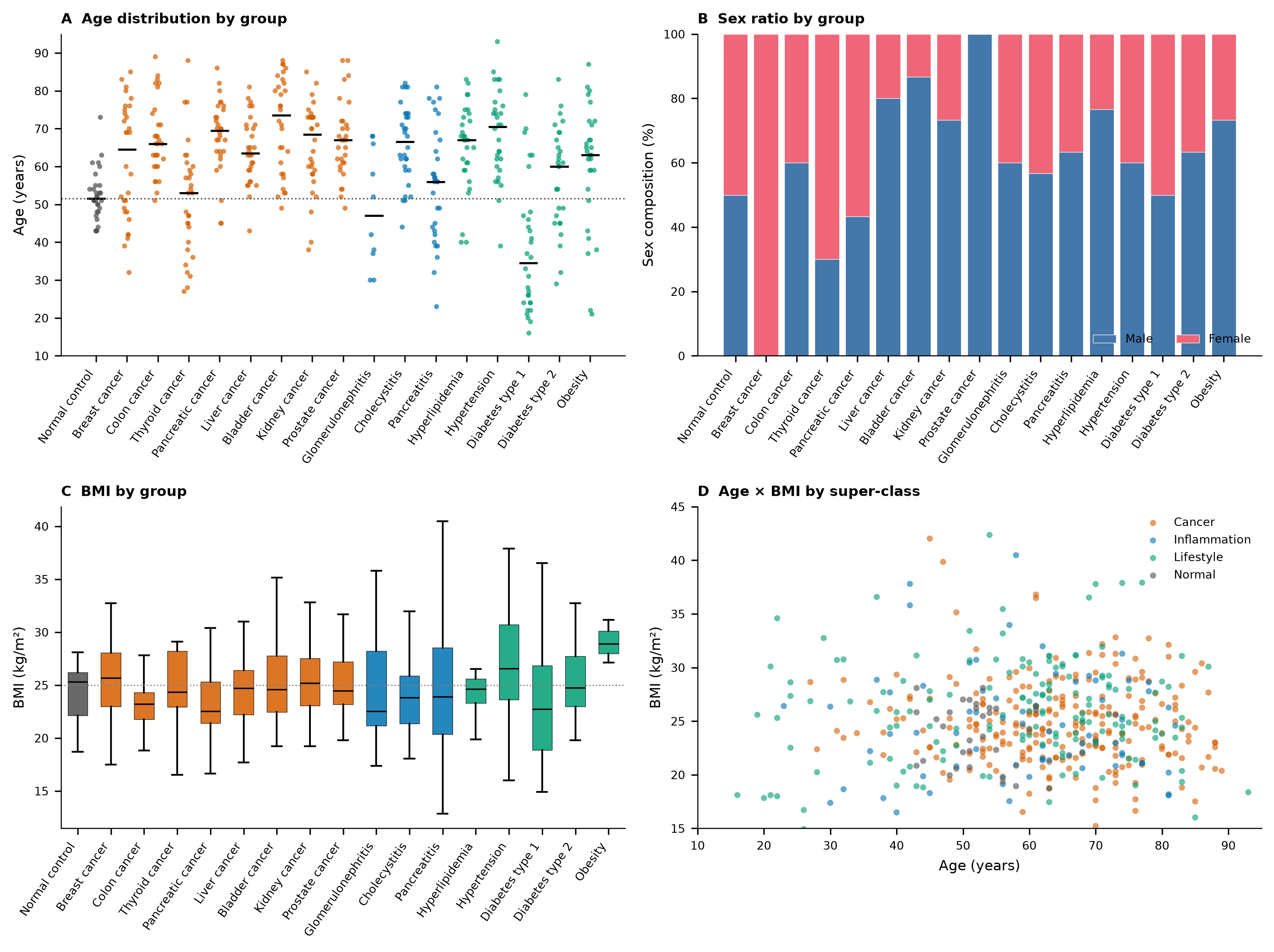

**Fig. S1 | Cohort demographics.** (A) Age distribution by group (points, individual participants; horizontal bars, group means; dotted line, normal-control mean of 52.4 years). (B) Sex composition (percentage male versus female) by group. (C) Body-mass-index distribution by group (boxes, median and IQR; whiskers, 1.5× IQR; dotted line, normal-control mean of 24.2 kg/m²). (D) Age versus BMI for all participants, coloured by super-class. The normal-control group is demographically representative of, rather than younger or leaner than, the disease groups; some individual groups differ in age or sex (for example the younger type-1-diabetes group and the female-predominant breast-cancer group), motivating the age/sex adjustment applied throughout the analysis.

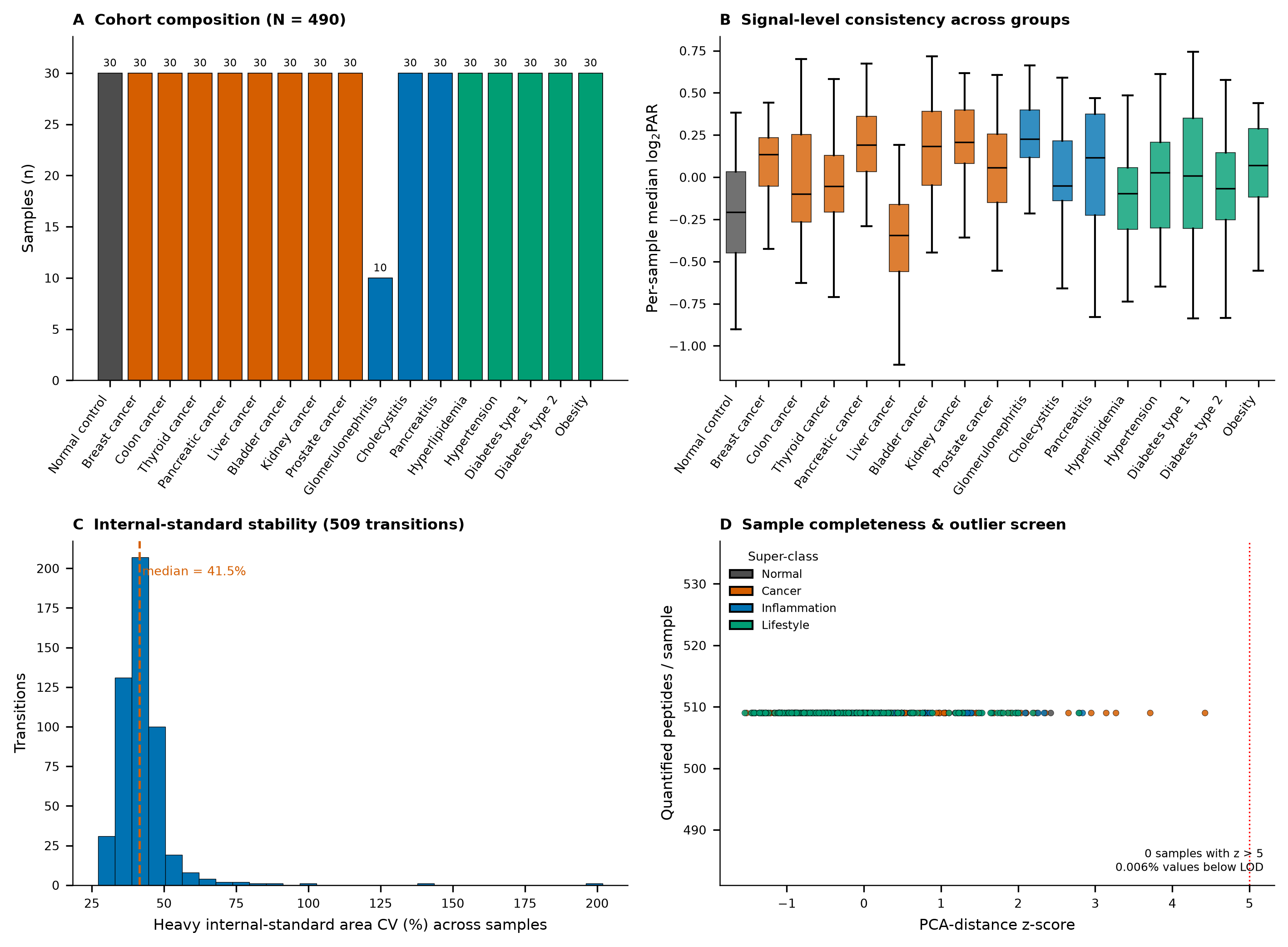

**Fig. S2 | Quality control.** (A) Cohort composition by group (N = 490), coloured by super-class; all groups contain 30 participants except glomerulonephritis (n = 10). (B) Per-group distributions of the per-sample median log₂ PAR, summarising overall signal level across groups before normalisation; boxes show the median and interquartile range and whiskers extend to 1.5× IQR. (C) Distribution of the coefficient of variation (CV) of the heavy internal-standard peak areas across samples for the 509 usable transitions; the median CV is 41.5% (dashed line), consistent with crude-plasma targeted assays. (D) Sample completeness and multivariate-outlier screen: quantified peptides per sample versus PCA-distance z-score, coloured by super-class. No sample exceeded z > 5 (dotted line) and only 0.006% of values fell below the limit of detection, indicating a clean, near-complete data matrix.

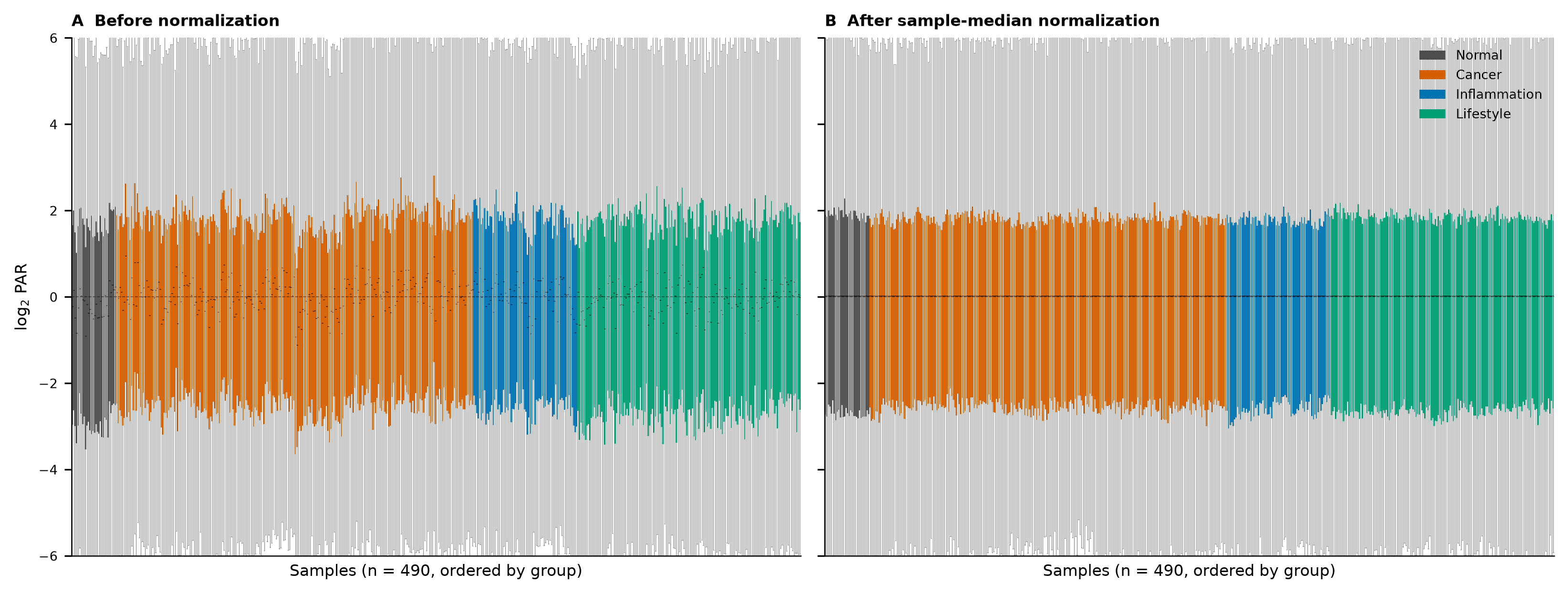

**Fig. S3 | Per-sample median normalisation.** Per-sample log₂ PAR distributions for all 490 samples, ordered by group and coloured by super-class, (A) before and (B) after per-sample median centring. Normalisation removes a modest per-sample loading offset—compressing the between-sample spread of per-sample medians from a standard deviation of 0.356 to zero—while preserving the within-sample dynamic range and the biological differences between proteins. Grey bars show the full per-protein spread within each sample and coloured bars mark the central interquartile region. Run-order drift, tested by Spearman correlation of per-sample median abundance against replicate index, was nominal in four of 17 groups and did not survive multiple-testing correction.

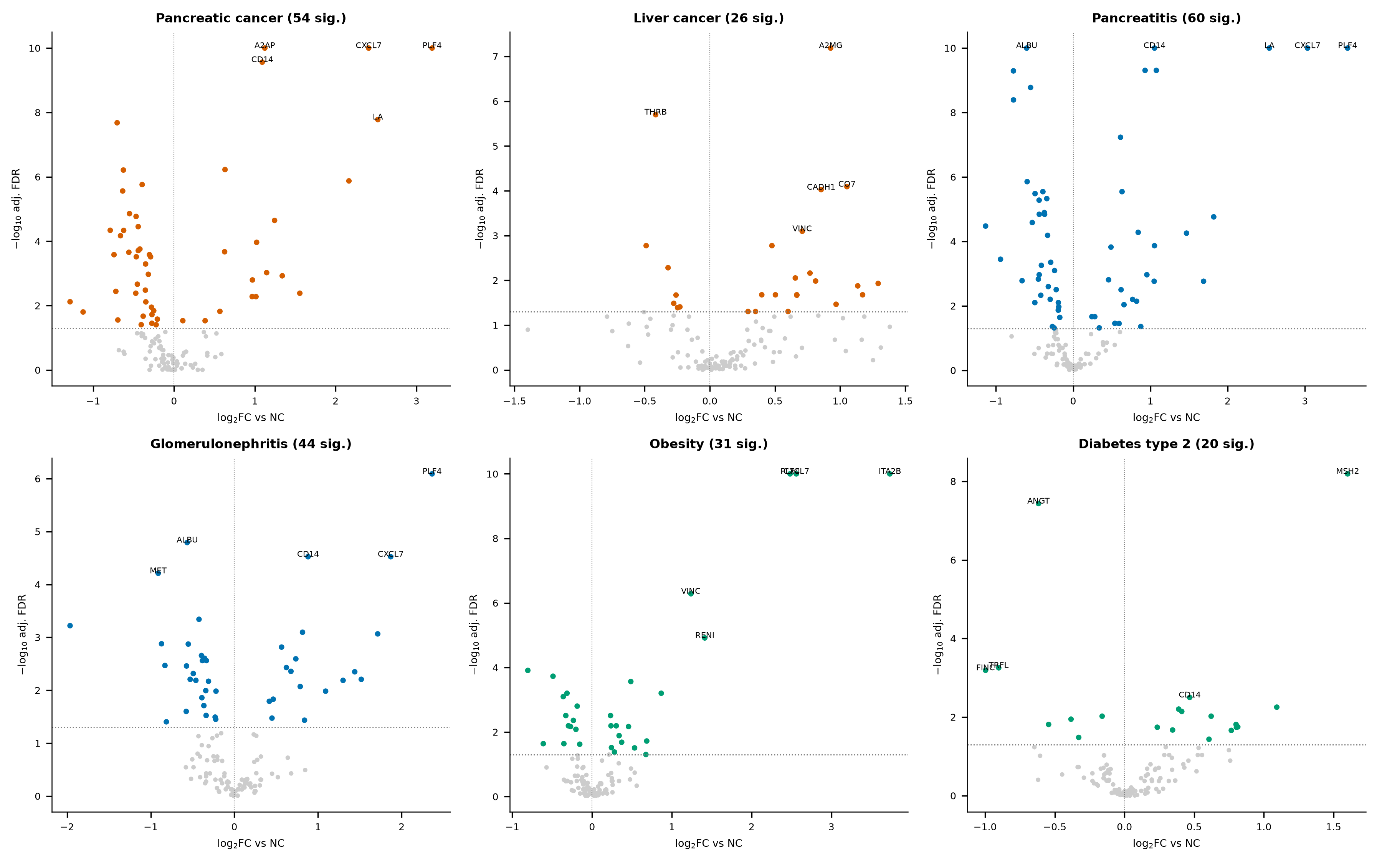

**Fig. S4 | Volcano plots of age/sex-adjusted differential abundance.** Age/sex-adjusted differential abundance for six representative diseases (pancreatic cancer, liver cancer, pancreatitis, glomerulonephritis, obesity, and type-2 diabetes), plotting −log₁₀ adjusted FDR against log₂ fold-change versus normal control. Coloured points are significant at FDR < 0.05 (the number of significant proteins is given in each panel title) and grey points are non-significant. Selected markers are labelled, including the broadly elevated CXCL7, PLF4, and ITA2B and disease-specific changes such as albumin (ALBU) in glomerulonephritis and ANGT and RENI in metabolic disease. Horizontal dotted lines mark the FDR = 0.05 threshold and vertical dotted lines mark zero fold-change.

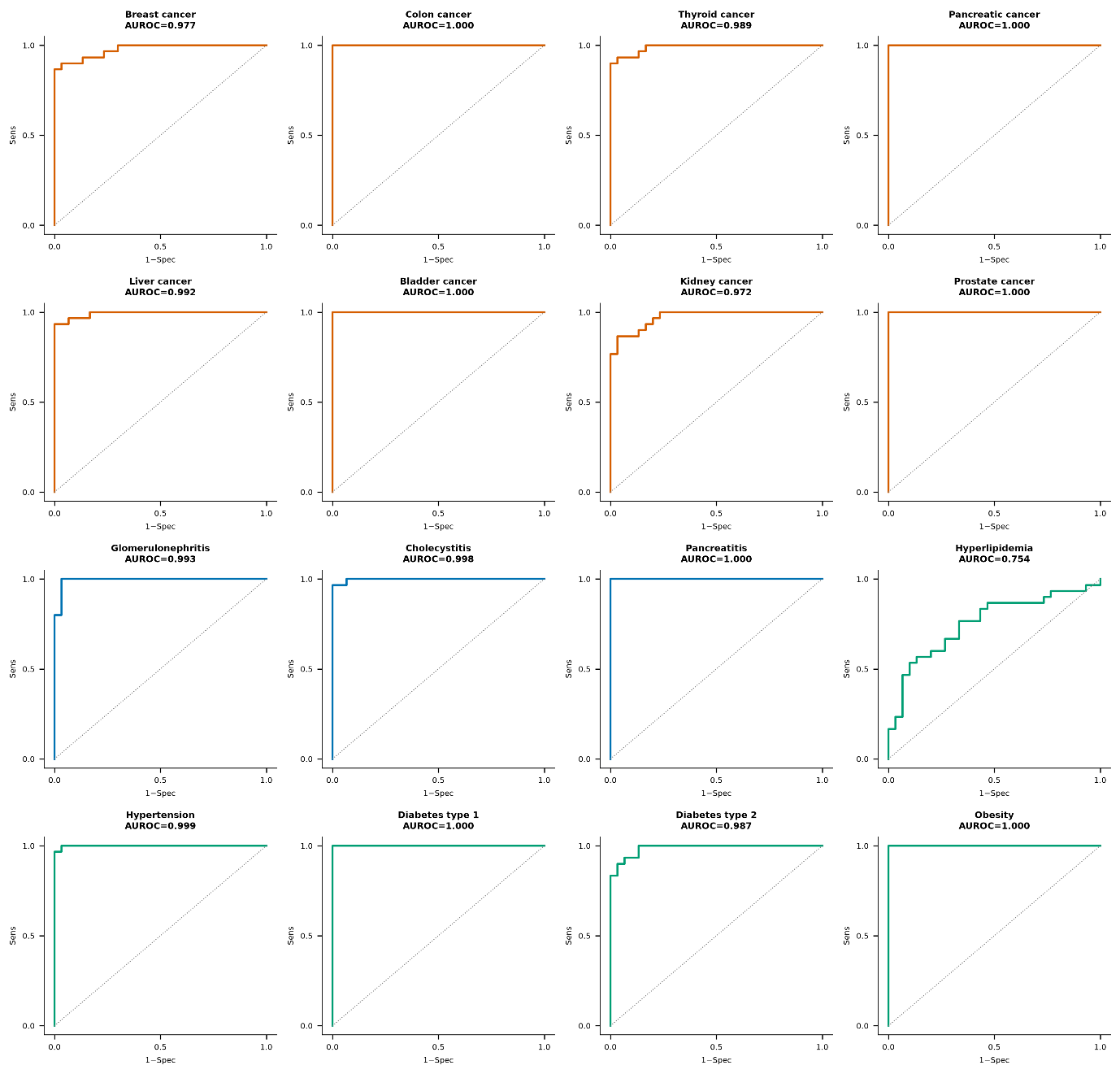

**Fig. S5 | Per-disease detection ROC curves.** Five-fold cross-validated receiver-operating-characteristic curves for each of the 16 diseases versus matched normal controls, with per-disease AUROC annotated and curves coloured by super-class; the diagonal marks chance. Most diseases show near-square curves (AUROC ≥ 0.97), several reaching 1.000, whereas hyperlipidaemia (AUROC 0.754) approaches the diagonal, consistent with a proteome signature that overlaps substantially with matched controls.

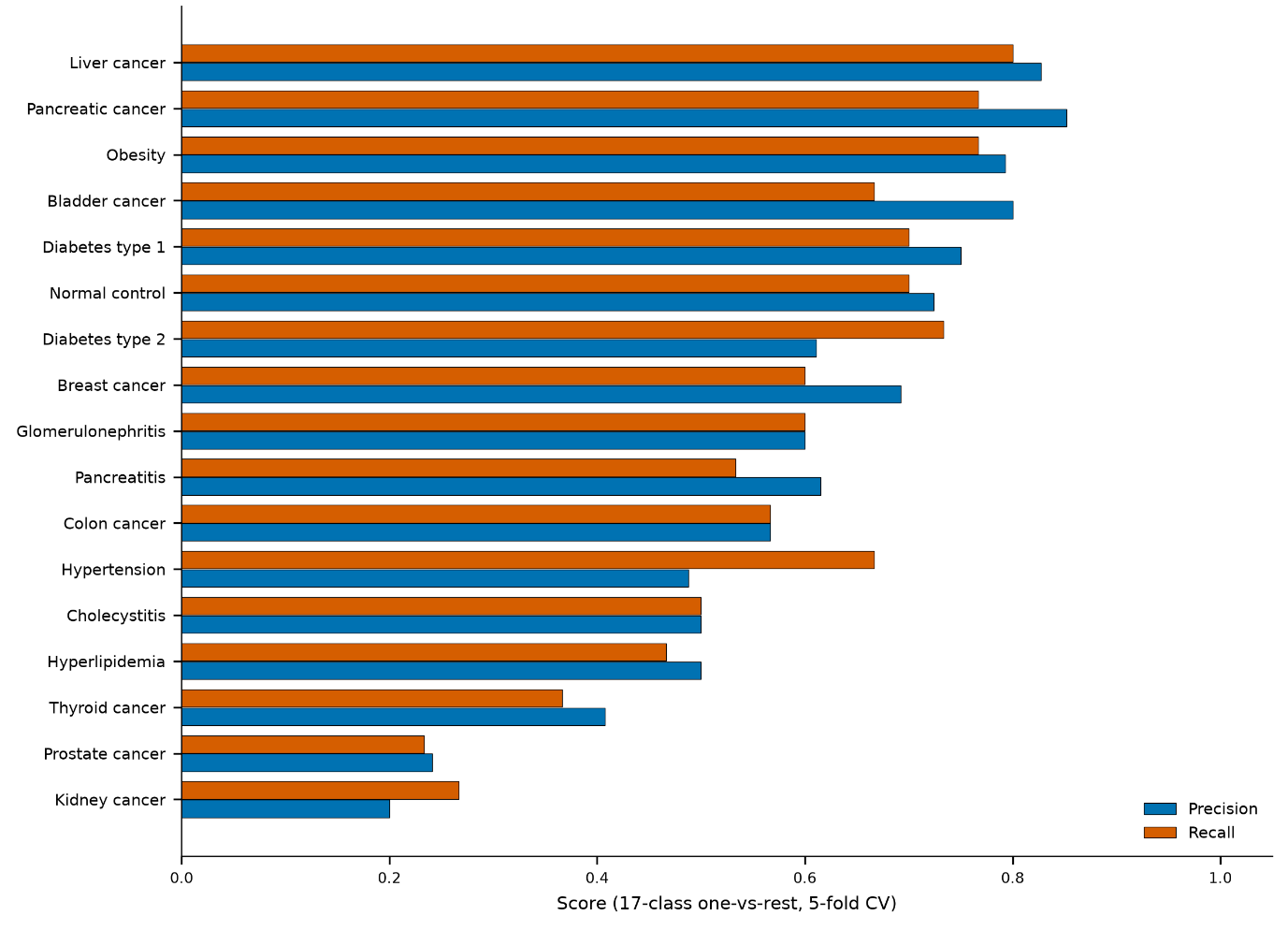

**Fig. S6 | Multi-class identification: per-class precision and recall.** Per-class precision (blue) and recall (orange) from the 17-class multinomial identification model (one-vs-rest, five-fold cross-validation), ordered by performance. Classes with distinctive proteomes (liver and pancreatic cancer, obesity, type-1 diabetes) achieve both high precision and recall, whereas overlapping conditions (kidney and prostate cancer, hyperlipidaemia) show lower values.

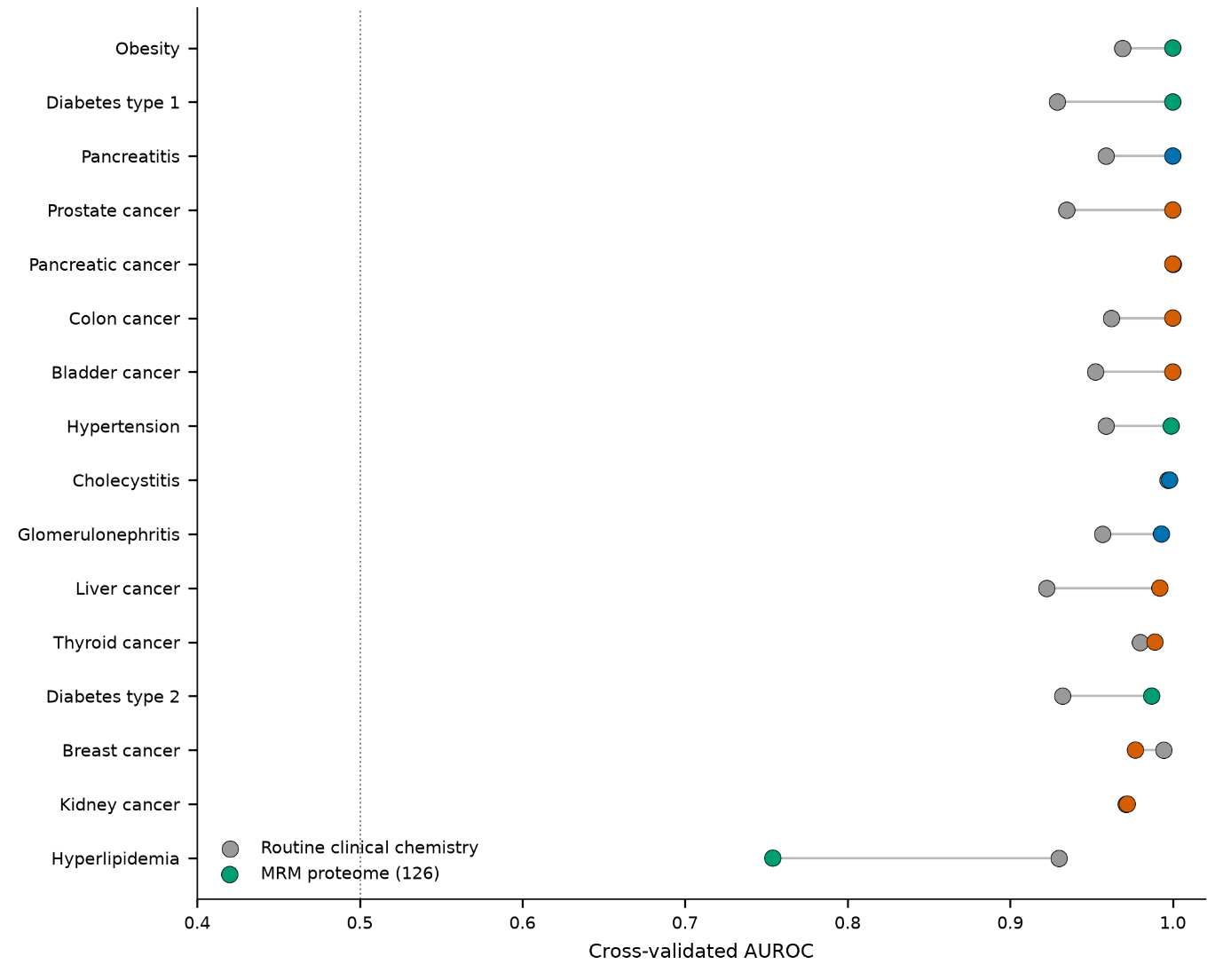

**Fig. S7 | MRM proteome versus routine clinical chemistry for per-disease detection.** Per-disease cross-validated detection AUROC obtained from the 126-protein MRM proteome (coloured points, by super-class) compared with that obtained from routine clinical-chemistry and haematology values alone (grey points), connected per disease; the vertical dotted line marks chance (0.5). Routine laboratory values discriminate most diseases at near-ceiling AUROC—implausible for genuine early screening—indicating that the cohort carries case-control batch or selection structure that any sufficiently rich feature set can exploit. This benchmark is the central evidence that the within-cohort accuracies are optimistic upper bounds rather than deployable screening accuracies.

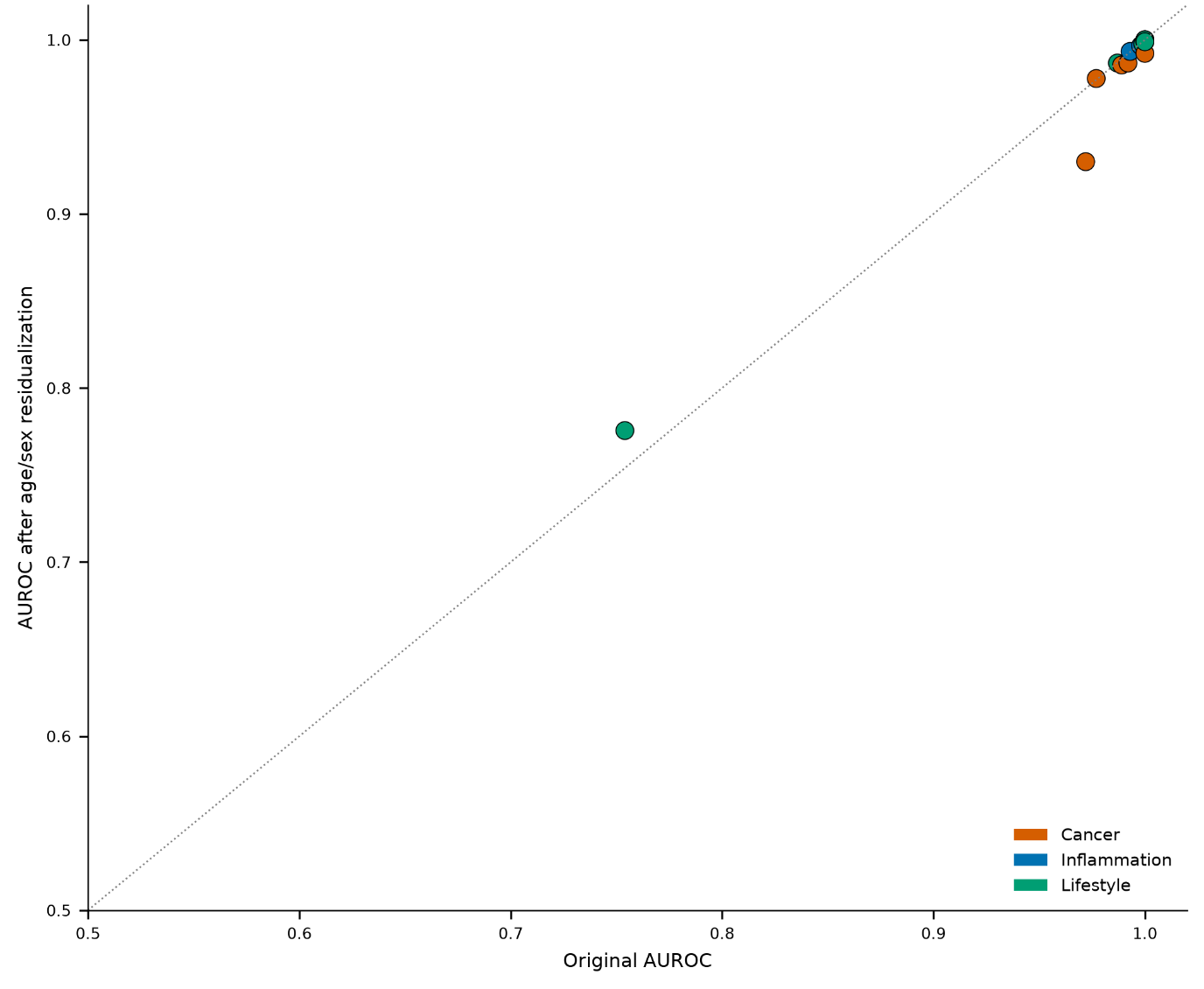

**Fig. S8 | Screening performance after removing age/sex variance.** Per-disease detection AUROC before (x-axis) versus after (y-axis) residualising every protein on age and sex, coloured by super-class; the dotted line is the identity. Points lie almost exactly on the diagonal (mean AUROC change 0.003), showing that the detection signal is not carried by age or sex and is therefore not a demographic artefact. The single markedly lower point corresponds to hyperlipidaemia, whose weaker underlying signal is the most sensitive to residualisation.

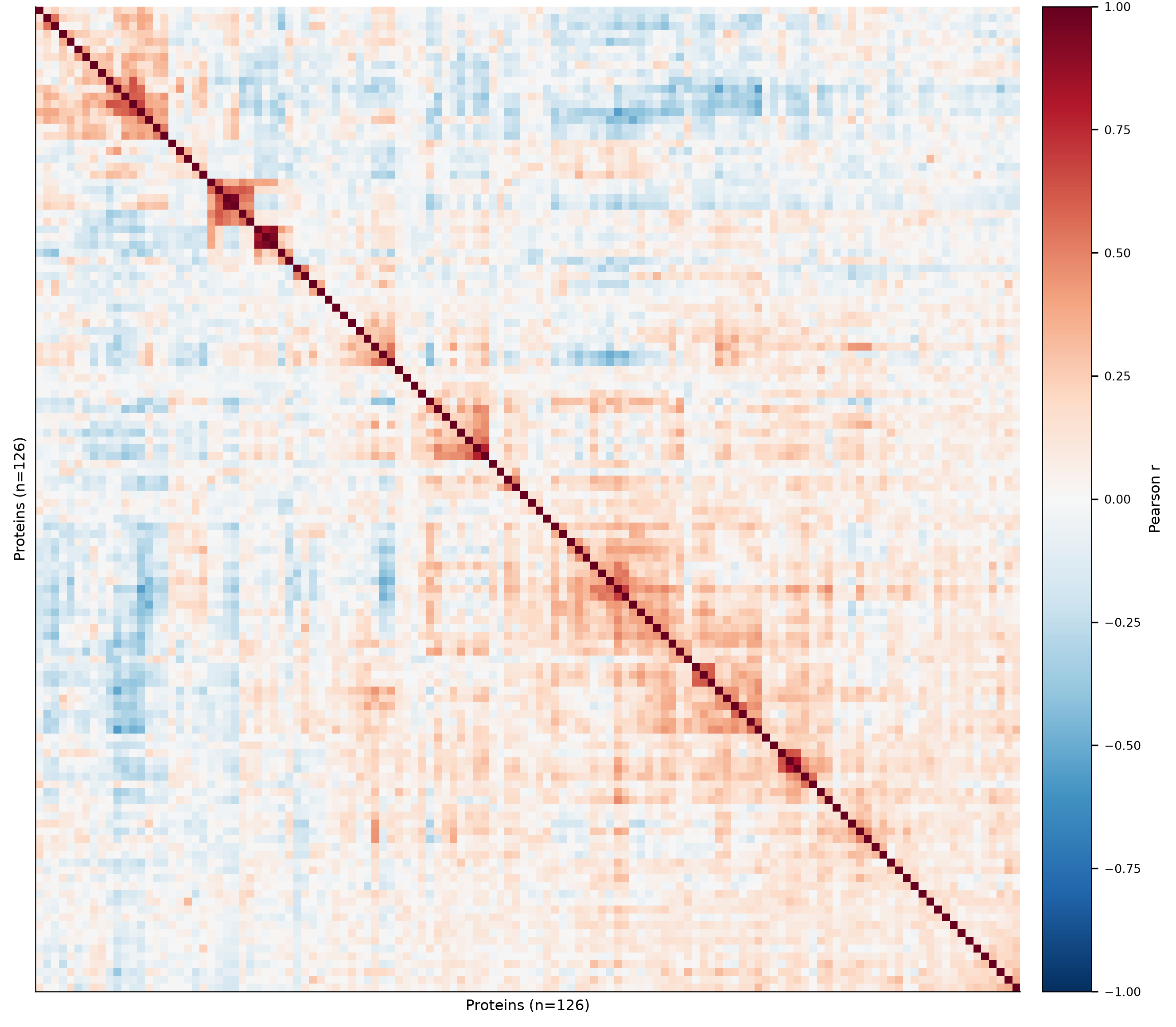

**Fig. S9 | Protein co-abundance structure.** Hierarchically ordered 126 × 126 matrix of pairwise Pearson correlations between protein abundances across all 490 samples (red, positive; blue, negative). A few small blocks of co-regulated proteins appear along the diagonal, but off-diagonal correlations are low overall (median absolute inter-protein |r| = 0.085), indicating that the panel—and the minimal 20-protein subset drawn from it—is largely non-redundant, with each protein contributing substantially independent information.

**Supplementary Data**

**Table S1.** Per-group demographics and routine clinical chemistry/haematology, with tests versus normal controls.

**Table S2.** Full protein-level differential-abundance results for all 16 diseases (Mann–Whitney and age/sex-adjusted, with fold-changes and FDR).

**Table S3.** Multi-class per-class metrics and per-disease detection performance.

**Table S4.** 20-protein consensus panel, per-disease top-five markers, and the panel-size performance curve.

Processed data matrices. peptide_matrix_norm.parquet and protein_matrix_norm.parquet (normalised abundance); peptide_PAR_matrix.parquet and protein_PAR_matrix.parquet (pre-normalisation PAR); harmonized_metadata.csv (clinical metadata); shap_importance.csv, panel_size_curve.csv, and differential_abundance_all.csv (analysis outputs).
